# Shifts in diversification rates and host jump frequencies shaped the diversity of host range among *Sclerotiniaceae* fungal plant pathogens

**DOI:** 10.1101/229930

**Authors:** Olivier Navaud, Adelin Barbacci, Andrew Taylor, John P. Clarkson, Sylvain Raffaele

## Abstract

The range of hosts that a parasite can infect in nature is a trait determined by its own evolutionary history and that of its potential hosts. However, knowledge on host range diversity and evolution at the family level is often lacking. Here, we investigate host range variation and diversification trends within the *Sclerotiniaceae*, a family of Ascomycete fungi. Using a phylogenetic framework, we associate diversification rates, the frequency of host jump events, and host range variation during the evolution of this family. Variations in diversification rate during the evolution of the Sclerotiniaceae define three major macro-evolutionary regimes with contrasted proportions of species infecting a broad range of hosts. Host-parasite co-phylogenetic analyses pointed towards parasite radiation on distant hosts long after host speciation (host jump or duplication events) as the dominant mode of association with plants in the *Sclerotiniaceae*. The intermediate macro-evolutionary regime showed a low diversification rate, high frequency of duplication events, and the highest proportion of broad host range species. Consistent with previous reports on oomycete parasites, our findings suggest that host jump and radiation, possibly combined with low speciation rates, could associate with the emergence of generalist pathogens. These results have important implications for our understanding of fungal parasites evolution and are of particular relevance for the durable management of disease epidemics.

## INTRODUCTION

The host range of a parasite has a central influence on the emergence and spread of disease (Woolhouse & Gowtage-Sequeria, 2005). There is a clear demarcation between specialist parasites that can only infect one or a few closely related host species, and generalists that can infect more than a hundred unrelated host species (Woolhouse et al., 2001, Barrett et al., 2009). Host specialization, when lineages evolve to infect a narrower range of hosts than related lineages, is a frequent occurrence in living systems and can be driven by a parasite sharing habitat with only a limited number of potential hosts. There are also clear examples of parasite adaptations that restrict the use of co-occuring potential hosts. For instance, the red rust fungal pathogen *Coleosporium ipomoeae* infects fewer *Ipomoea* species from its native community than when inoculated to nonnative communities, implying that evolution within local communities has narrowed pathogen host range (Chappell & Rausher, 2016). Some isolates of the rice blast fungus *Magnaporthe oryzae* are able to infect *Oryza sativa* japonica varieties but not indica varieties co-occuring in the Yuanyang area of China, because their genome harbors numerous avirulence effector genes (Liao et al., 2016). It has been proposed that specialization results from trade-offs between traits needed to infect a wide range of hosts (Futuyma & Moreno, 1988, Joshi & Thompson, 1995). It is frequently associated with the loss of traits that are not required to infect a particular host, such as the loss of lipid synthesis in parasitoid wasps (Visser et al., 2010), and the loss of secondary metabolism and carbohydrate active enzymes in powdery mildew fungal pathogens (Spanu et al., 2010). Specialization is sometimes considered as an evolutionary dead end (Moran, 1988), since gene losses are often irreversible and may lead specialist lineages "down a blind alley" that limits transitions back to generalism (Haldane, 1951, Day et al., 2016). There is nevertheless evidence for transitions from specialist to generalist parasitism (Johnson et al., 2009, Hu et al., 2014). For instance in plant pathogens, *Pseudoperonospora cubensis* differs from other downy mildew oomycete pathogens in that it is able to infect a wide range of Cucurbits (Thines & Choi, 2015). For many parasite lineages however, knowledge on host range diversity and evolution at the macro-evolutionary level is lacking.

*Sclerotiniaceae* is a family of Ascomycete fungi of the class Leotiomycetes which includes numerous plant parasites. Among the most studied are the grey mould pathogen *Botrytis cinerea*, considered to be one of the 10 most devastating plant pathogens (Dean et al., 2012), and the white and stem mold pathogen *Sclerotinia sclerotiorum*. Both are economically important pathogens in agriculture that infect hundreds of host plant species (Bolton et al., 2006, Mbengue et al., 2016). The *Sclerotiniaceae* family also includes host specialist parasites such as *Ciborinia camelliae* that causes flower blight on *Camellia* (Denton-Giles et al., 2013), *Sclerotinia glacialis* that specifically infects *Ranunculus glacialis* (Graf & Schumacher, 1995), and *Monilinia oxycocci* causing the cottonball disease on cranberry (McManus et al., 1999). Other species from the *Sclerotiniaceae* have intermediate host range (tens of plant species) such as *Sclerotinia trifoliorum*, *S. subarctica* and *S. borealis* (Farr & Rossman, 2016, Clarkson et al., 2010). While *S. sclerotiorum* and *B. cinerea* are considered as typical necrotrophic pathogens, rapidly killing host cells to cause disease, the *Sclerotiniaceae* include species with diverse lifestyles. For instance, the poplar pathogen *Ciborinia whetzelii* and several *Myriosclerotinia* species are biotrophs that can live as endophytes (Schumacher & Kohn, 1985, Andrew et al., 2012), while *Coprotinia minutula* is coprophilous (Elliott, 1967). How this remarkable diversity evolved remains elusive. To gain insights into this question, knowledge of phylogenetic relationships and host range diversity at the macro-evolutionary level is needed. Specifically, ancestral state reconstruction can provide insights into how fungal host range has changed over time. The relationship between host range and diversification rates is an active area of research in insect ecology (Hamm & Fordyce, 2015, Hardy & Otto, 2014). For instance, a positive correlation was reported between species richness in the butterfly family *Nymphalidae* and the diversity of plants they feed upon (Janz et al., 2006). However in contrast, some studies found a negative relationship between host-plant breadth and diversification rate (Hardy & Otto, 2014). Transitions between feeding strategies generally appeared to be associated with shifts in insect diversification rates (Janz & Nylin, 2008, Hardy & Otto, 2014). Estimating fungal species divergence times in the *Sclerotiniaceae* will allow testing of whether biological diversification is related to host range variation.

Host range is a trait determined not only by the evolutionary history of a parasite, but also by that of its potential hosts (Poulin & Keeney, 2008). Accounting for host association patterns should therefore prove useful to understand host range evolution in the *Sclerotiniaceae*. Leotiomycete class diverged less than 200 million years ago (Prieto & Wedin, 2013, Beimforde et al., 2014) and likely radiated with the diversification of flowering plants (Smith et al., 2010). Molecular phylogenetic studies distinguished *Myriosclerotinia*, *Sclerotinia sensu stricto*, *Botrytis* and *Botryotinia*, and *Monilinia sensu stricto* as monophyletic clades within the *Sclerotiniaceae* family (Holst-Jensen et al., 1998, Holst-Jensen et al., 2004). A phylogenetic analysis of *Monilinia* species suggested that co-speciation with host plants was the dominant pattern in this clade (Holst-Jensen et al., 1997). By contrast, there was no evidence for co-speciation with host plants in a phylogenetic analysis of *Botrytis* species (Staats et al., 2005). *Botrytis* species were thus proposed to have evolved through host jumps to unrelated host plants followed by adaptation to their new hosts (Staats et al., 2005, Dong et al., 2015). Host-parasite co-phylogenetic analyses are required to test whether variations in the frequency of host jumps may have impacted on host range variation in the *Sclerotinaceae*.

In this study, we performed a phylogenetic analysis on 105 *Sclerotiniaceae* species to reveal multiple independent shifts and expansions of host range. We show that three macro-evolutionary regimes with distinct diversification rates and dominant host-association patterns have shaped the diversity of the *Sclerotiniaceae*, and lead to contrasted proportions of broad host range species. Specifically, we highlight an increased emergence of broad host range parasites during the transition between macro-evolutionary regimes dominated by distinct patterns of host-pathogen association. These results suggest that reduced diversification rates and high host-jump frequency could associate with the emergence of generalist pathogens.

## MATERIALS AND METHODS

### Taxon and host range data selection

We used all 105 *Sclerotiniaceae* species for which at least *ITS* marker sequence data were available in the Genbank database. As outgroups we selected 56 *Rutstroemiaceae* species and 39 representative species of Leotiomycetes and Sordariomycetes for a total of 200 species. Host range data were obtained from (Boland & Hall, 1994, Melzer et al., 1997), Index Fungorum (http://www.indexfungorum.org), the SMML fungus-host distribution database (Farr & Rossman, 2016) and references therein. In total, we retrieved 7101 fungus-host association records that did not show strong geographic or crop/wild species bias (**Supplementary Figure 1**). Calibrated trees deciphering the relationships between the host families based on 7 gene regions (18S rDNA, 26S rDNA, ITS, *matK*, *rbcL*, *atpB*, and *trnL-F*) were extracted from (Qian & Zhang, 2014) and updated with (Hedges et al., 2015). The tree of host families used for cophylogenetic analyses is provided as Supplementary File 1.

### Initial phylogenetic analysis

*ITS* sequences were aligned using MAFFT version 7 (Katoh & Standley, 2013). The alignment was manually adjusted to minimize possible homoplasic positions using Seaview 4 (Gouy et al., 2010). We retained gaps shorter than 41 positions present in a maximum of 39 sequences, with ungapped blocks being at least five characters long with a maximum of 5% ambiguous nucleotides per position. Unalignable and autapomorphic regions were excluded from the analysis, yielding an alignment with 797 informative sites (**Supplementary File 2**). Maximum-likelihood Phylogeny was inferred with PhyML 3 (Guindon et al., 2010) with Smart Model Selection which permits an automatic substitution model selection supporting the general time-reversible model with gamma distribution using 6 substitution rate categories (GTR+G6) model as the best fit. Statistical branch support was inferred with the SH-like approximate likelihood ratio test (SH-aLRT) (Anisimova et al., 2011) and bootstrap analysis with 100 replicates (**Supplementary File 3 and 4**). The topology of the tree was confirmed by three additional methods. First, we used a neighbor-joining approach in FastME 2.0 (Lefort et al., 2015) with the LogDet substitution model and tree refinement by subtree pruning and regrafting (**Supplementary File 5**). Second, we used the parsimony ratchet approach implemented in the phangorn package in R (Schliep, 2010) (**Supplementary File 6**). Third, we used Bayesian analysis in MrBayes 3.2.6 (Ronquist et al., 2012) using the GTR substitution model with gamma-distributed rate variation across sites and a proportion of invariable sites in 100 million MCMC generations, sampling parameters every 1,000, removing 10% of the tree files as burn-in. The resulting trees were edited with Figtree (available at http://tree.bio.ed.ac.uk/software/figtree/).

### Ancestral state reconstruction

To infer possible ancestral hosts, we used Reconstruct Ancestral State in Phylogenies 3.1 (RASP) (Yu et al., 2015). We used the S-DIVA (Statistical-Dispersal Vicariance Analysis), S-DEC (Statistical Dispersal– Extinction–Cladogenesis model), BBM (Bayesian Binary MCMC) and BayArea methods to verify congruence between the methods and assess the robustness of their output, as recommended by (Yu et al., 2015). We report the results of the S-DIVA analysis which is considered the best adapted for host-parasite association analyses (Yu et al., 2015, Razo-Mendivil & De Leon, 2011). To implement S-DIVA, we delimited host groups as follows: Vitales (A), Asterids (B), Campanuliids (C), Commelinids (D), coprophilous (E), core Eudicots (F), Eudicots (G), Fabids (H), Polypodiidae (I), Lamiids (J), Magnoloidae (K), Malvids (L), Monocotyledones (M), Pinidae (N), allowing a maximum of 4 groups at each node. Among these groups, the latest divergence is that of Malvids and Fabids, estimated around 82.8 and 127.2 Mya (Clarke et al., 2011) and predates the emergence of *Sclerotiniaceae* estimated at ˜69.7 Mya in this work, we therefore did not restrict associations in the ancestral reconstruction analysis. To account for incomplete sampling in this analysis, ancestral state reconstruction was computed for every plant group by the re-rooting method (Yang et al., 1995) under an entity-relationship (ER) model available in phytools (Revell, 2012) and derived from the ape library in R (Paradis et al., 2004). This method based on maximum likelihood, computed for every nodes of the *Sclerotiniaceae* phylogeny the probability to infect a given group of hosts. We considered plant groups as hosts when the probability of the ancestral state was >50%. For every plant group, up to 10% of terminal nodes were pruned randomly 100 times and the ancestral state reconstructed on pruned trees. For each plant group, we then extracted the variation in the age of the most ancestral inclusion into the *Sclerotiniaceae* host range (**Supplementary Figure 2**). For all plant groups, this variation was not significantly different from 0, indicating that ancestral state reconstruction was robust to tree pruning.

### Divergence dating analyses

We used a Bayesian approach to construct a chronogram with absolute times with the program BEAST 1.8.2. As no fossil is known in this fungal group, we use Sordariomycetes-Leotiomycetes divergence time to calibrate the tree (^~^300 Mya, Beimforde et al., 2014). Using our initial phylogeny as a constraint, the partition file was prepared with the BEAUti application of the BEAST package. Considering that our dataset includes distantly related taxa, subsequent analyses were carried out using an uncorrelated relaxed clock model with lognormal distribution of rates. We used a uniform birth-death model with incomplete species sampling as prior on node age, following (Beimforde et al., 2014) to account for incomplete sampling in our phylogeny. Analyses were run three times for 100 million generations, sampling parameters every 5,000 generations, assessing convergence and sufficient chain mixing using Tracer 1.6 (Drummond et al., 2012a). We removed 20% of trees as burn-in, and the remaining trees were combined using LogCombiner (BEAST package), and summarized as maximum clade credibility (MCC) trees using TreeAnnotator within the BEAST package. The trees were edited in FigTree. The R packages *ape* (Paradis et al., 2004) and *phytools* (Revell, 2012) were used for subsequent analyses.

### Diversification analysis

We used the section containing all 105 *Sclerotiniaceae* species of the chronogram with absolute times generated by BEAST to perform diversification analyses with three different methods. We first used Bayesian Analysis of Macroevolutionary Mixtures (BAMM) version 2.5 (Rabosky et al., 2013, Rabosky et al., 2014), an application designed to account for variation in evolutionary rates over time and among lineages. The priors were set using the setBAMMPriors command in the BAMMtools R-package (Rabosky et al., 2014) . We ran four parallel Markov chains for 5,000,000 generations and sampled for every 1000 trees. The output and subsequent analyses were conducted with BAMMtools. We discarded the first 10% of the results and checked for convergence and the effective sample size (ESS) using the *coda* R-package. Lineage-through-time plots with extant and extinct lineages were computed using the phytool package in R (Revell, 2012) with the drop.extinct=FALSE parameter. The exact number of *Sclerotiniaceae* species is currently unknown and no intentional bias was introduced in data collection (**Supplementary Figure 1**). To detect shifts in diversification rates taking into account incomplete sampling of the *Sclerotiniaceae* diversity and tree uncertainty, we used MEDUSA, a stepwise approach based upon the Akaike information criterion (AIC) (Alfaro et al., 2009, Drummond et al., 2012b). For this analysis, we used the birth and death (bd) model, allowed rate shifts both at stem and nodes, used the AIC as a statistical criterion with initial r=0.05 and epsilon=0.5. Sampling of the fungal diversity is typically estimated around 5% (Hawksworth & Luecking, 2017). We used random sampling rates between 5 and 100% for each individual species in 100 MEDUSA bootstrap replicates to estimate the impact of sampling richness. Next, we randomly pruned 1 to 10% of species from the tree in 100 MEDUSA bootstrap replicates to estimate the impact of tree completeness. To control for the impact of divergence time estimates on the diversification analysis, we altered all branching times in the tree randomly by -15% to +15% in 100 MEDUSA bootstrap replicates (**Supplementary Figure 3**). Finally, we used RPANDA which includes model-free comparative methods for evolutionary analyses (Morlon et al., 2016). The model-free approach in RPANDA compares phylogenetic tree shapes based on spectral graph theory (Lewitus & Morlon, 2015). It constructs the modified Laplacian graph of a phylogenetic tree, a matrix with eigenvalues reflecting the connectivity of the tree. The density profile of eigenvalues (**Supplementary Figure 4A**) provides information on the entire tree structure. The algorithm next used k-medioids clustering to identify eight modalities in the phylogenetic tree of the *Sclerotiniaceae* (**Supplementary Figure 4B-C**). A post-hoc test comparing Bayesian Information Criterion (BIC) values for randomly bifurcated trees (BIC_random_) with that of the tree of the *Sclerotiniaceae* (BIC_Sclerotiniaceae_) supported the eight modalities (BIC_random_/BIC_Sclerotiniaceae_ = 10.34, well above the significance threshold of 4.0). We obtained significant BIC ratios with two or more modalities, supporting the existence of at least two macro-evolutionary regimes in the *Sclerotiniaceae.*

### Co-phylogeny analysis

To detect potential codivergence patterns, we tested for co-phylogeny between fungal species and host family trees with CoRe-PA 0.5.1 (Merkle et al., 2010), PACo (Balbuena et al., 2013) and Jane 4 (Conow et al., 2010) as these programs can accommodate pathogens with multiple hosts. The only 3 species which were not able to interact with an angiosperm host (saprotroph or only described on Gymnosperms) were excluded from the analysis (namely: *Elliottinia kerneri*, *Coprotinia minutula* and *Stromatinia cryptomeriae*). We used a full set of 263 host-pathogen associations and a simplified set of 121 associations minimizing the number of host families involved to control for the impact of sampling bias (**Table 1**). We also performed the analyses independently for each macro-evolutionary regime. To estimate cost parameters in CoRe-PA, we ran a first cophylogeny reconstruction with cost values calculated automatically using the simplex method on 1000 random cycles, with all host switches permitted, direct host switch, and full host switch permitted (other parameters set to CoRe-PA default). The best reconstruction had a quality score of 1.348 with total costs 37.076 and calculated costs of 0.0135 for cospeciation, 0.1446 for sorting, 0.2501 for duplication and 0.5918 for host switch. With these costs and parameters, we then computed reconstruction with 1000 random associations to test for the robustness of the reconstructions. Next, we used 100,000 permutations of the host-pathogen association matrix in PACo to test for the overall congruence between host and pathogen trees. To test for the contribution of each association on the global fit, we performed taxon jackknifing. We used one-tailed z tests to compare squared residuals distribution for each host-pathogen link with the median of squared residuals for the whole tree, and infer likely co-evolutionary and host shift links at p<0.01. We used optimal costs calculated by CoRe-PA to run co-phylogeny reconstruction in Jane 4 Solve Mode with 50 generations and a population size of 1,000. Statistical tests were performed using 100 random tip mapping with 20 generations and population size of 100. For the analysis of each macro-evolutionary regime in Jane 4, we used simplified host-pathogen association set to reduce the amount of Losses and Failure to Diverge predicted and facilitate comparisons with CoRe-PA and PACo results. We visualized the interactions with TreeMap3 (Charleston & Robertson, 2002), using the untangling function to improve the layout.

**TABLE 1.** Results of the cophylogeny analyses under CoRe-PA optimized cost settings using various methods and host association sets. ^a^ refers to the set of host association tested for cophylogeny: ‘Full set’ corresponds to the complete list of all *plant-Sclerotiniaceae* associations; ‘Simplified’ corresponds to a reduced set covering the whole *Sclerotiniaceae* family; ‘G1’, ‘G2’,and ‘G3’ corresponds to associations involving *Sclerotiniaceae* species from macro-evolutionary regime G1, G2 or G3 only. ^b^ standard deviation correspond to frequencies calculated for 1000 reconstruction with randomized host-parasite associations. ^c^ In PACo taxon jackknifing, associations that contributed significantly and positively to cophylogeny were classified as ‘cospeciation’, significantly and negatively as ‘host switch’ and associations with no significant contribution to cophylogeny are classified as either Sorting/loss or Duplication. ^d^ values correspond to CoRe-Pa total reconstruction costs, p-value of the observed host-parasite association matrix m^2^ in 100,000 permutations with PACo, or the p-value of the observed reconstruction cost in 100 random tip mappings in Jane 4. FD, failure to diverge.

| Method | Host associations<sup>a</sup> | Event frequency (% of host associations) |  |  |  |  | Cost / p-val<sup>d</sup> |
| --- | --- | --- | --- | --- | --- | --- | --- |
|  |  | Cospeciation | Sorting/Loss | Duplication | Host switch | FD |  |
| **CoRe-PA<sup>b</sup>** | **Full set (263)** | **15.82 ± 2.34** | **15.90 ± 4.80** | **34.35 ± 3.35** | **33.94 ± 3.49** |  | **38.21 ± 0.79** |
|  | **Simplified (121)** | **9.63 ± 2.63** | **11.09 ± 4.97** | **40.20 ± 3.25** | **39.07 ± 4.00** |  | **40.44 ± 0.81** |
|  | G1 only (68) | 15.75 ± 3.19 | 16.32 ± 7.35 | 37.78 ± 4.67 | 30.14 ± 5.23 |  | 13.95 ± 0.44 |
|  | G2 only (130) | 17.03 ± 3.57 | 16.21 ± 7.58 | 33.16 ± 5.27 | 33.60 ± 5.63 |  | 14.57 ± 0.47 |
|  | G3 only (65) | 11.78 ± 5.17 | 14.38 ± 8.69 | 32.86 ± 6.75 | 40.98 ± 6.87 |  | 8.95 ± 0.34 |
| **PACo<sup>c</sup>** | **Full set (263)** | **20.91** | **48.29** |  | **30.79** |  | **p<1.0<sup>-05</sup>** |
|  | **Simplified (121)** | **15.70** | **57.02** |  | **27.27** |  | **p<1.0<sup>-05</sup>** |
|  | G1 only (68) | 27.94 | 55.88 |  | 16.18 |  | p=2.0<sup>-05</sup> |
|  | G2 only (130) | 6.15 | 71.54 |  | 22.31 |  | p= 0.00045 |
|  | G3 only (65) | 0.00 | 66.15 |  | 33.85 |  | p= 0.00607 |
| **Jane 4** | **Full set (263)** | **0.86** | **67.73** | **8.00** | **3.57** | **19.83** | **1291 (p<0.01)** |
|  | **Simplified (121)** | **2.90** | **65.22** | **14.78** | **11.59** | **5.51** | **719.47 (p<0.01)** |
|  | G1 simple (43) | 1.39 | 41.67 | 25.00 | 26.39 | 5.56 | 206.98 (p<0.01) |
|  | G2 simple (54) | 2.28 | 75.80 | 11.87 | 3.65 | 6.39 | 374.08 (p<0.01) |
|  | G3 simple (24) | 9.68 | 25.81 | 19.35 | 41.94 | 3.23 | 105.41 (p=0.13) |

## RESULTS

### Multiple independent shifts and expansions of host range in the evolution of the *Sclerotiniaceae*

To document the extant diversity in the *Sclerotiniaceae* fungi, we collected information on the host range of 105 species in this family. For comparison purposes, we also analyzed the host range of 56 species from the *Rutstroemiaceae* family, a sister group of the *Sclerotiniaceae* (**Figure 1A, Table 1**). To reduce biases that may arise due to missing infection reports, we analyzed host range at the family level instead of genus or species level. We found one species in the *Sclerotiniaceae* (*Coprotinia minutula*) and two species in the *Rutstroemiaceae* (*Rutstroemia cunicularia* and *Rutstroemia cuniculi*) reported as non-pathogenic to plants that are coprophilous (Elliott, 1967). Most species in the *Sclerotiniaceae* are reported necrotrophic pathogens, such as *Botrytis*, *Monilinia* and *Sclerotinia* species. Exceptions include *Ciborinia whetzelii* which was reported as an obligate biotroph of poplar (Andrew et al., 2012) and *Myriosclerotinia* species that specifically infect monocot families as facultative biotrophs or symptomless endophytes (Andrew et al., 2012). The most frequently parasitized plant group was Fabids (by 47 *Sclerotiniaceae* species and 26 *Rutstroemiaceae* species), a group of the Rosids (Eudicots) including notably cultivated plants from the Fabales (legumes) and Rosales (rose, strawberry, apple…) orders. Plants from the order Vitales (e.g. grapevine), from the Magnoliids, and from the *Polypodiidae* (ferns) were colonized by *Sclerotiniaceae* but not *Rutstroemiaceae* fungi.

We next used the internal transcribed spacer *(ITS)* region of rDNA sequences to construct a phylogenetic tree of the 105 *Sclerotiniaceae* and 56 *Rutstroemiaceae* species (**Figure 1B, Sup. File 1-7**). This marker is the only one available for all species and is currently considered as the universal marker for taxonomic use (Schoch et al., 2012). Maximum likelihood, neighbor joining, parsimony and Bayesian analyses yielded convergent tree topologies. The tree topologies confirmed the familial classification recognized in previous phylogenies (Holst-Jensen et al., 1997, Holst-Jensen et al., 1998, Andrew et al., 2012), identifying a clearly supported *Botrytis* genus including 23 species; a *Myriosclerotinia* genus including 8 species; a *Sclerotinia* genus *sensu stricto* including 4 *Sclerotinia*; and a *Monilinia* genus including 16 *Monilinia* species and 3 other species. *Botrytis calthae* and *Amphobotrys ricini* were the only two species with imprecise positions in maximum likelihood reconstructions, due to the *ITS* marker being insufficient to infer complete phylogenetic placement (Staats et al., 2005, Lorenzini & Zapparoli, 2016).

**Figure 1.**
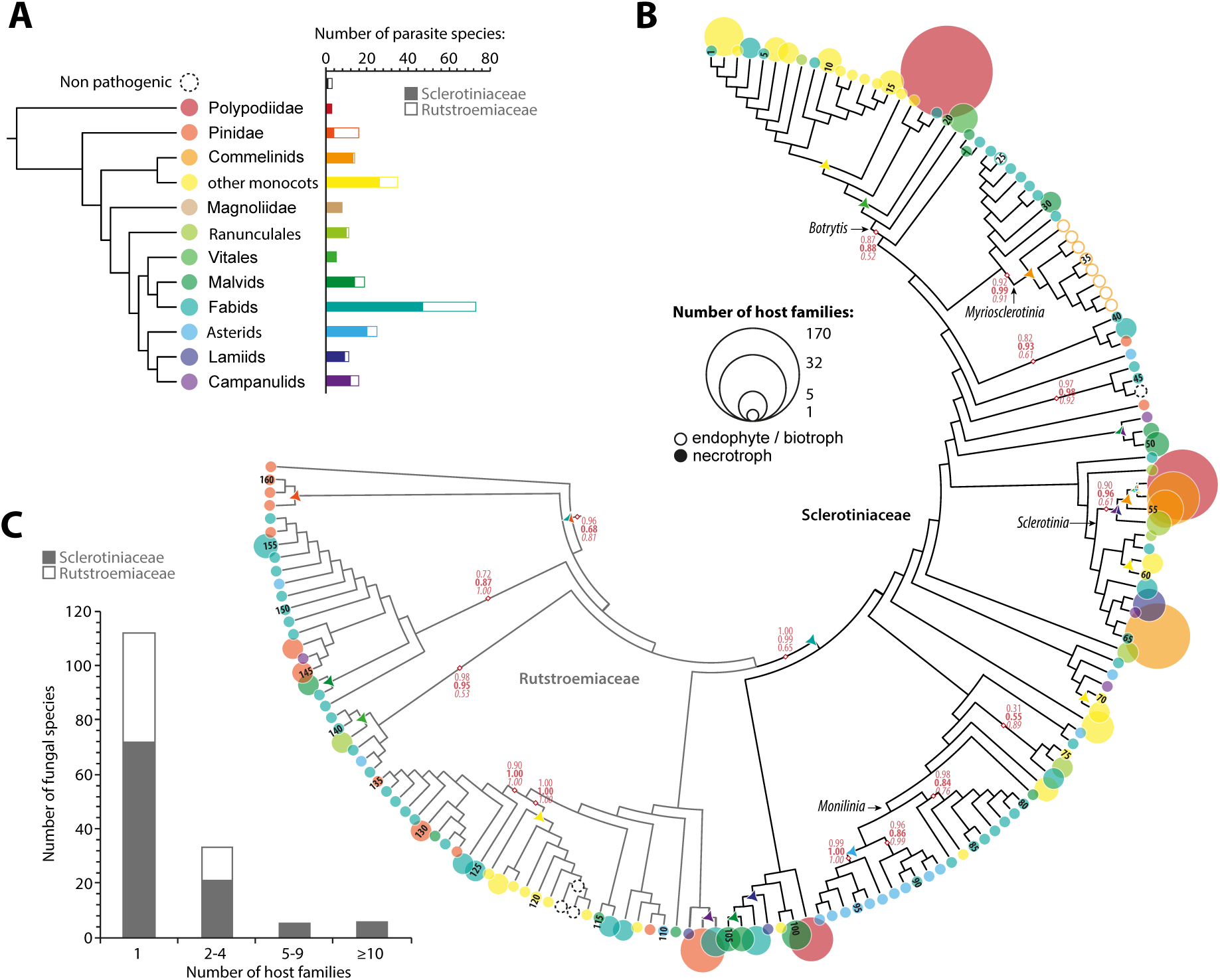
Multiple independent shifts and expansions of host range in the evolution of the *Sclerotiniaceae*. (**A**)Distribution of plant hosts of parasites from the *Sclerotiniaceae* and *Rutstroemiaceae* fungi.( **B**) Maximum likelihood *ITS* phylogeny of 105 *Sclerotiniaceae* and 56 *Rutstroemiaceae* species obtained by showing host range information and ancestral host reconstruction. Host range is shown as circles at the tips of branches, sized according to the number of host families and colored as in (A) according to the earliest diverging plant group in host range. Numbers at the tips of branches refer to species listed in Supplementary Table 1. Branch support indicated in light red for major clades corresponds to SH-aLRT (regular), bootstrap (bold) and Bayesian posterior probabilities (italics). Reconstructed ancestral host is shown as triangles at intermediate nodes when a change compared to the previous node is predicted. Endophytes and biotrophic parasites are shown with empty circles. (**C**) Distribution of *Sclerotiniaceae* and *Rutstroemiaceae* species according to their number of host families.

We used RASP (Yu et al., 2015) and phytools (Revell, 2012) to reconstruct the ancestral host range across the phylogeny (**Figure 1B, Supplementary Figure 1, Supplementary Figure 5**). This identified Fabids as the most likely ancestral hosts of the *Sclerotiniaceae* family (relative probability of 100% in S-DIVA and S-DEC analyses). Random tree pruning in phytools indicated that this result is robust to sampling biases. Over 48% of the *Sclerotiniaceae* species are pathogens of host plants that evolved prior to the divergence of the Fabids, suggesting numerous host jumps in this family of parasites. Notably, a jump to Malvids and then to Monocots was identify at the base of the *Botrytis* genus (87% and 85% probability in S-DIVA respectively), a jump to Commelinids occurred in the *Myriosclerotinia* genus (91% probability in S-DIVA), a jump to the Ranunculales occurred at the base of the *Sclerotinia* genus (76% probability in S-DIVA); a jump to Asterids was found at the base of a major group of *Monilinia* (89% probability in S-DIVA).

A total of 73 *Sclerotiniaceae* species (69.5%) and 41 (73.2%) *Rutstroemiaceae* infected a single host family (**Figure 1C, Supplementary Table 1**). *Moellerodiscus lentus* was the only *Rutstroemiaceae* species infecting hosts from more than 5 plant families whereas eleven *Sclerotiniaceae* species (10.4%) exhibited this trait, including *Botrytis cinerea*, *Sclerotinia sclerotiorum*, *Sclerotinia minor* and *Grovesinia pyramidalis*, each of which colonizes plants from more than 30 families. Each of these species belong to a clearly distinct phylogenetic group with a majority of species infecting a single host family. This may result from radiation following host jumps (Choi & Thines, 2015) and suggests that the ability to colonize a broad range of plant was acquired multiple times independently through the evolution of the *Sclerotiniaceae*.

### Two major diversification rate shifts in the evolution of the *Sclerotiniaceae*

To test for a relationship between host range variation and biological diversification in the *Sclerotiniaceae*, we estimated divergence times for the fungal family *Sclerotiniaceae* using the *ITS* marker in a Bayesian framework. A calibrated maximum clade credibility chronogram from these analyses is shown in **Figure 2A** (and **Supplementary Figure 6**). This showed that *Sclerotiniaceae* fungi shared a most recent common ancestor between 33.9 and 103.5 million years ago (Mya), with a mean age of 69.7 Mya. The divergence of *Botrytis pseudocinerea,* estimated from the *ITS* dataset, occurred ca. 3.35-17.8 Mya (mean age 9.8 Mya) and is similar to a previous estimate of 7-18 Mya (Walker et al., 2011).

**Figure 2.**
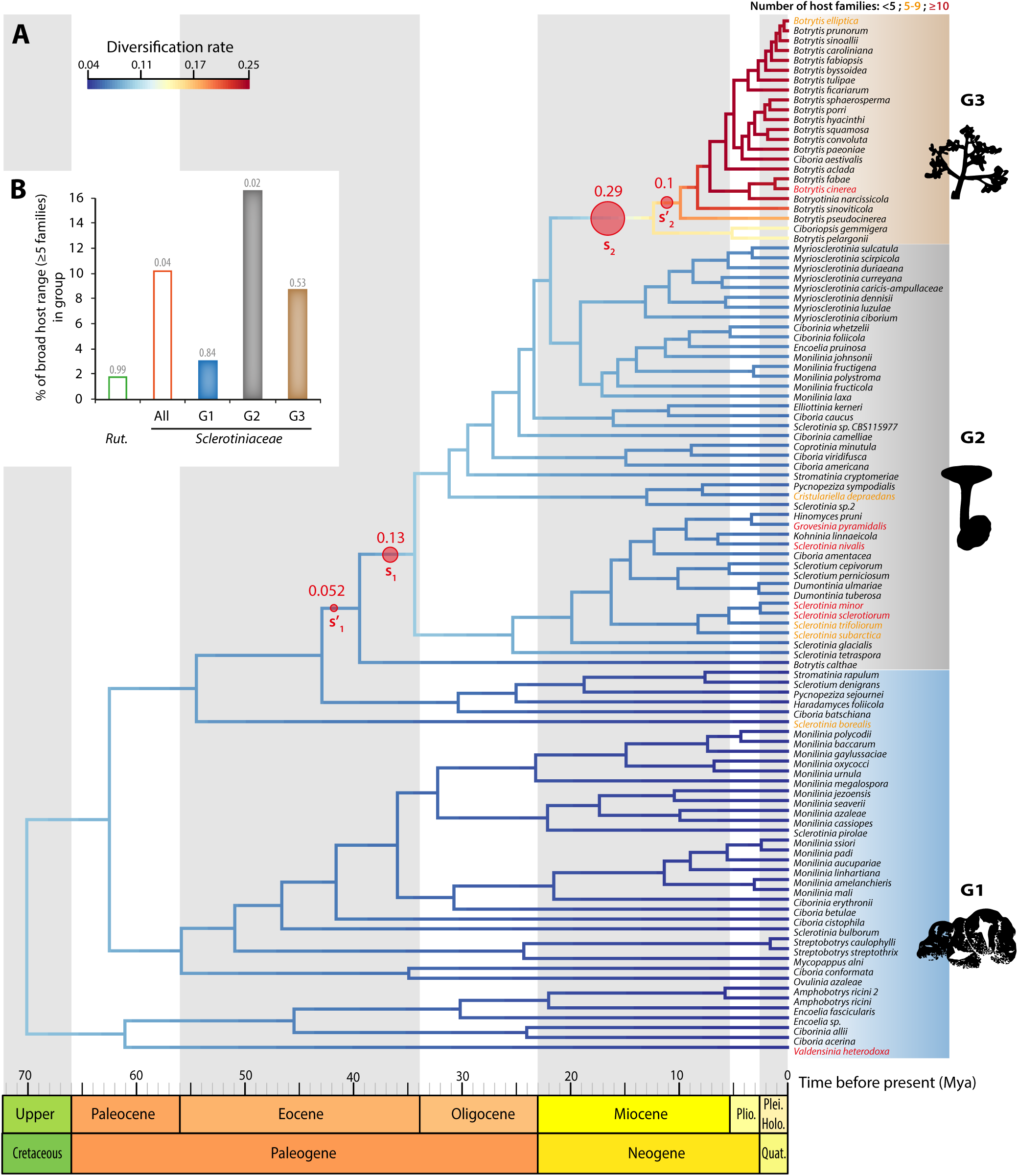
Two major diversification rate shifts in the evolution of the *Sclerotiniaceae.* (A) Dated ITS-based species tree for the *Sclerotiniaceae* with diversification rate estimates. The divergence times correspond to the mean posterior estimate of their age in millions of years calculated with BEAST. Branches of the tree are color-coded according to diversification rates determined with BAMM. Major rate shifts identified in BAMM are shown as red circles, noted s1, s′1, s2 and s′2 and labeled with the posterior distribution in the 95% credible set of macro-evolutionary shift configurations. Species names are shown in black if host range includes less than 5 plant families, in yellow for 5 to 9 plant families and in red for 10 or more plant families. Diversification rate shifts define three macro-evolutionary regimes noted G1, G2 and G3 and boxed in blue, grey and brown respectively. (**B**) Distribution of broad host range (5 or more host families) parasites in *Rutstroemiaceae, Sclerotiniaceae* and under each macro-evolutionary regime of the *Sclerotiniaceae*. P-values calculated by random permutations of host ranges along the tree are indicated above bars. Holo., Holocene; Mya, Million years ago; Plei., Pleistocene; Plio., Pliocene; Quat. Quaternary; Rut., *Rutstroemiaceae*.

Using this time-calibrated phylogeny, we calculated speciation and diversification rates using three different methods, to control for the limits of individual methods (Rabosky et al., 2017). First, we used the BAMM framework, which implements a Metropolis Coupled Markov chain Monte Carlo method to calculate diversification rates along lineages (Rabosky, 2014) (**Figure 2A, Supplementary Figure 7A-C**). The analysis identified two significant rate shifts (s_1_ and s_2_, accounting for > 30% of the posterior distribution in the 95% credible set of macro-evolutionary shift configurations) each immediately adjacent to a minor rate shift (s_1_′ and s_2_′ respectively; ≤10% of the posterior distribution) within two different clades (**Figure 2A**). Each shift affects a specific clade and is not detected coincidentally across the whole phylogeny. The earliest shift occurred between 34.2 and 42.7 Mya. It resulted in an increase of instantaneous diversification rate from ^~^0.05 lineage/million year in *Monilinia* and related clades to ^~^0.08 lineage/million year in *Sclerotinia* and derived clades. The second shift occurred between 9.8 and 21.7 Mya. It resulted in a further increase of instantaneous diversification to >0.15 lineage/million year in *Botrytis* lineage. These two shifts defined three distinct macro-evolutionary regimes in the *Sclerotiniaceae*. Second, to determine how speciation rates changed over time, we computed lineage-through-time plots for the *Sclerotiniaceae* tree and 1000 trees in which we altered all branching times randomly by -15% to +15% to control for the sensitivity to divergence time estimates (**Figure 3A**). We observed an exponential trend of species accumulation consistent with a late burst of cladogenesis or early extinction (Pybus γ=0.284). Third, we used MEDUSA (Alfaro et al., 2009, Drummond et al., 2012b) to detect shifts in diversification rate in the *Sclerotiniaceae* phylogeny. MEDUSA reported two shifts, corresponding to shifts s_1_ and s_2_′ identified in BAMM (**Sup. Fig. 3D**). We used a bootstrap approach to test for the sensitivity of these shifts to sampling richness, tree completeness and dating accuracy (**Figure 3B, Supplementary Figure 3**). Shift s_2_′ was detected in 100% of the bootstraps. Shift s1 was the second most frequent shift detected in our bootstrap analysis although it appeared sensitive to sampling richness in particular. Estimates of speciation rates in MEDUSA were robust to sampling richness, tree completeness and dating accuracy (r^~^0.038 in G1, r^~^0.074 in G2 and r^~^0.31 in G3). These values were consistent with diversification rate estimated in BAMM for each regime (**Figure 3C**). Fourth, we used model-free analysis of branching patterns in phylogenetic trees with R-PANDA (Morlon et al., 2016). The spectrum of eigenvalues suggested eight modes of division in the phylogeny of the *Sclerotiniaceae* (**Supplementary Figure 4A-B**). Comparison to randomly bifurcating trees suggested that these modalities are significant (BIC ratio 0.09). The modalities identified by RPANDA k-means clustering algorithm overlapped with the three macro-evolutionary regimes identified previously (**Supplementary Figure 4C**). Overall, these analyses converged towards the identification of three macro-evolutionary regimes in the

**Figure 3.**
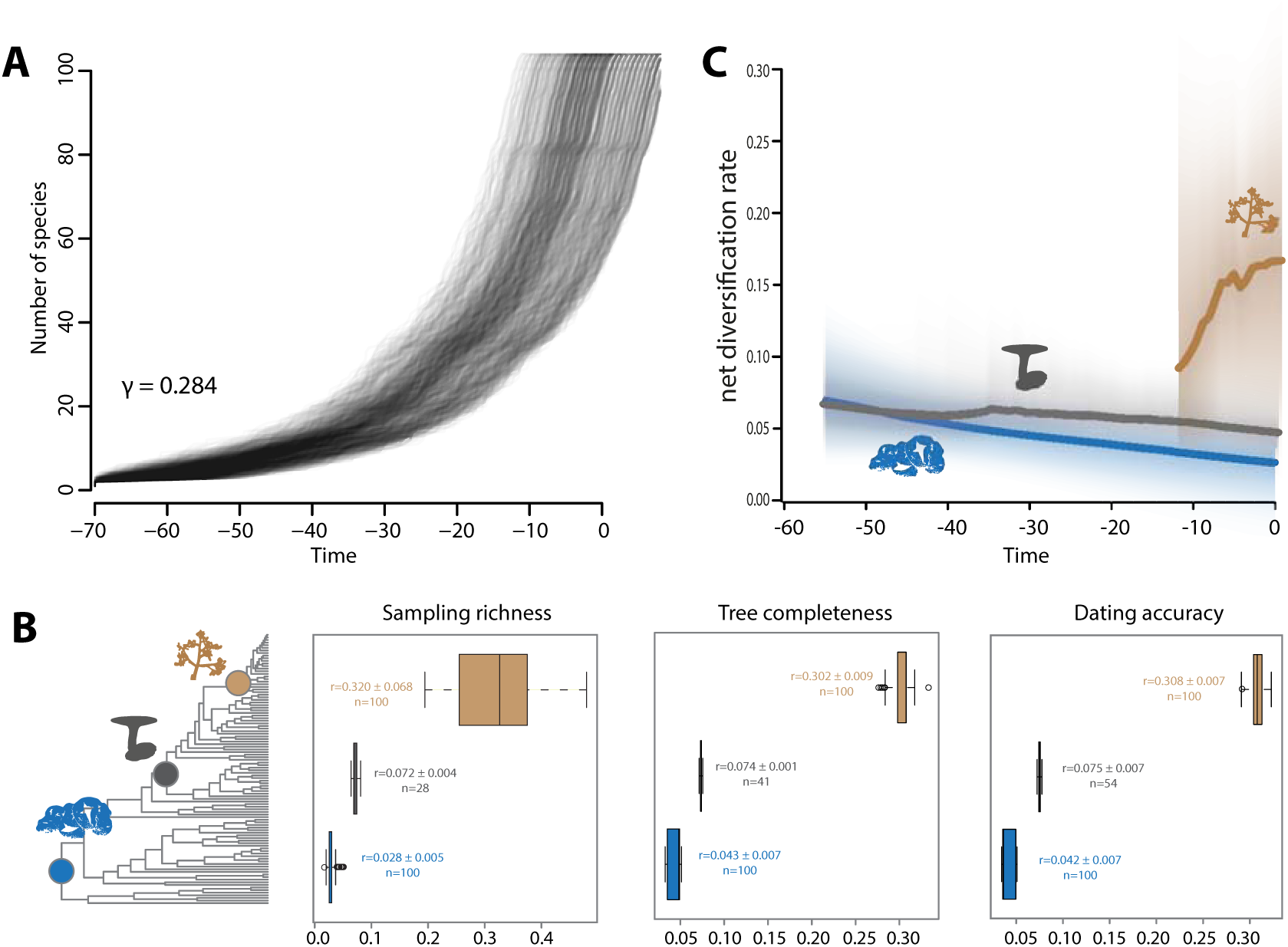
Robustness of diversification rate shifts identification in the Sclerotiniaceae phylogeny. (A)Lineage-through-time plots for the *Sclerotiniaceae* tree and 1000 trees in which branching times were altered randomly by -15% to +15% to control for the sensitivity to divergence time estimates. Pybus γ for the *Sclerotiniaceae* tree is provided. (**B**) Frequency (n) of diversification rate shift detection and diversification rate estimates (r) in a 100 MEDUSA bootstrap replicates in which sampling richness, tree completeness and divergence times were randomly altered. Labels indicate average diversification rate estimates (r) for each macro-evolutionary regime (blue for G1, grey for G2, brown for G3), with standard deviation of the mean for a 100 replicates. (**C**) Net diversification rates over time estimated by BAMM for each macro-evolutionary regime (blue for G1, grey for G2, brown for G3).

### Sclerotiniaceae

Species in regime G1 are early diverging and showed low diversification rates, they encompassed notably *Monilinia*, *Encoelia* and some *Ciboria* species. Regime G2 had intermediate diversification rates and encompassed notably *Myriosclerotinia* species and most *Sclerotinia* species. Regime G3 corresponding to *Botrytis* genus was the most recently diverged and presented the highest diversification rates.

We compared the proportion of broad host range (≥5 plant families) fungal species that emerged in the *Rutstroemiaceae*, in the *Sclerotiniaceae*, and under each the three macro-evolutionary regimes in the *Sclerotiniaceae* (**Figure 2B**). We randomly permuted host range values across the tree 10,000 times to estimate the p-values of these proportions occurring by chance. We counted one (1.8%, p=0.9959) broad host range species in the *Rutstroemiaceae* (*Moellerodiscus lentus*) and eleven (10.4%, p= 0.0359) in the *Sclerotiniaceae*. In the *Sclerotiniaceae*, regime G2 with intermediate diversification rates showed the highest proportion of broad host range species (16.2%, p=0.0163), followed by regime G3 (8.7% of broad host range species, p=0.537) and regime G1 (5.1% of broad host range species, p=0.8407). These analyses suggest that the evolutionary history of regime G2 could have favored the emergence of broad host range parasites in the *Sclerotiniaceae*.

### Diversification rate shifts associate with variations in rates of co-speciation, duplication and host jump in the*Sclerotiniaceae*

Within the last 50 Ma the world has experienced an overall decrease in mean temperatures but with important fluctuations that dramatically modified the global distribution of land plants (Zachos et al., 2001, Donoghue & Edwards, 2014, Nürk et al., 2015). We hypothesized that modifications in the distribution of plants could have an impact on host association patterns in *Sclerotiniaceae* parasites. To test this hypothesis, we performed host-parasite co-phylogeny reconstructions using CoRe-PA (Merkle et al., 2010), PACo (Balbuena et al., 2013) and Jane 4 (Conow et al., 2010) (**Table 1**). We found that hosts of *Sclerotiniaceae* fungi include ^~^1800 plant species. To consider this overall host diversity in a co-phylogenetic analysis, we used a tree including 56 plant families, a tree of 102 *Sclerotiniaceae* parasitic species, and 263 host-parasite interactions (**Sup. File 3**). The resulting tanglegram (**Figure 4A**) highlighted clear parasite duplications notably on *Ericaceae* and *Rosaceae* in the G1 group of the *Sclerotiniaceae* (*Monilinia* species). There was no obvious topological congruence between the plant tree and groups G2 and G3 of the *Sclerotiniaceae*. To take into account sampling bias, we performed co-phylogeny reconstructions using the full set of 263 host-parasite associations and a simplified set of 121 associations. Reconstructions were also performed independently on each of the three macro-evolutionary regimes. CoRe-PA classifies host-pathogen associations into (i) cospeciation, when speciation of host and pathogen occur simultaneously, (ii) duplication, when pathogen speciation occurs independently of host speciation, (iii) sorting or loss, when a pathogen remains associated with a single descendant host species after host speciation, and (iv) host switch (also designated as host jump) when a pathogen changes host independently of speciation events (Merkle et al., 2010). The CoRe-PA reconstructions indicated that duplications and host switch each represented ^~^34% of host associations from the full set of *host-Sclerotiniaceae* associations. Cospeciation and sorting each represented ^~^16% of host associations. The analysis of each macro-evolutionary regime indicated that G1 is characterized by high duplication and low host jump frequencies and G3 by low cospeciation and sorting frequencies and high host jump frequencies (**Figure 4B**). We next used Procrustean superimposition in PACo (Balbuena et al., 2013) to identify host-pathogen associations that contribute significantly to co-phylogeny between *Sclerotiniaceae* and host plant families. The PACo analysis suggested that overall, the *Sclerotiniaceae* lineages are not randomly associated with their host families (P<0.01). The PACo analysis also includes taxon jackknifing to test for the relative contribution of each host-pathogen association in the co-phylogeny pattern. In the full set of *host-Sclerotiniaceae* associations, ^~^21% contributed positively and significantly to cophylogeny, likely representing cospeciation events, while ^~^31% contributed negatively and significantly to cophylogeny, therefore likely representing host jump events (**Table 1**). Taxon jackknifing on individual macro-evolutionary regime revealed a high frequency of associations with positive contribution to cophylogeny (likely cospeciation) in G1, and a high frequency of associations with negative contribution to cophylogeny (likely host switch) in G3, in good agreement with the CoRe-PA analysis. In addition to the types of host-parasite association mentioned before, Jane 4 identifies ‘failure to diverge’ associations, corresponding to events when a host speciates and the parasite remains on both new host species (Conow et al., 2010). Jane 4 analysis for the whole *Sclerotiniaceae* family (simplified set of associations) identifies losses as the dominant form of host association (^~^65%), followed by duplications (^~^15%) and host jumps (^~^12%) while failure to diverge (^~^5.5%) and cospeciation (^~^3%) were rare events (**Table 1**). Consistent with CoRe-PA and PACo analyses, Jane 4 found the highest rate of host jumps in regime G3 (^~^42%). Unlike previous analyses, Jane 4 found the highest rate of cospeciation in G3 (^~^9.7%). G1 was characterized by a high duplication rate while G2 was characterized by a high rate of losses and failure to diverge events but the lowest host jump rate. Overall, our co-phylogeny analyses converged towards the conclusion that cospeciations represent a minor proportion of host associations in the *Sclerotiniaceae* and that the proportion of host jumps varied markedly between the three macro-evolutionary regimes, with G3 showing the highest host jump rate (between 33% and 42%).

**Figure 4.**
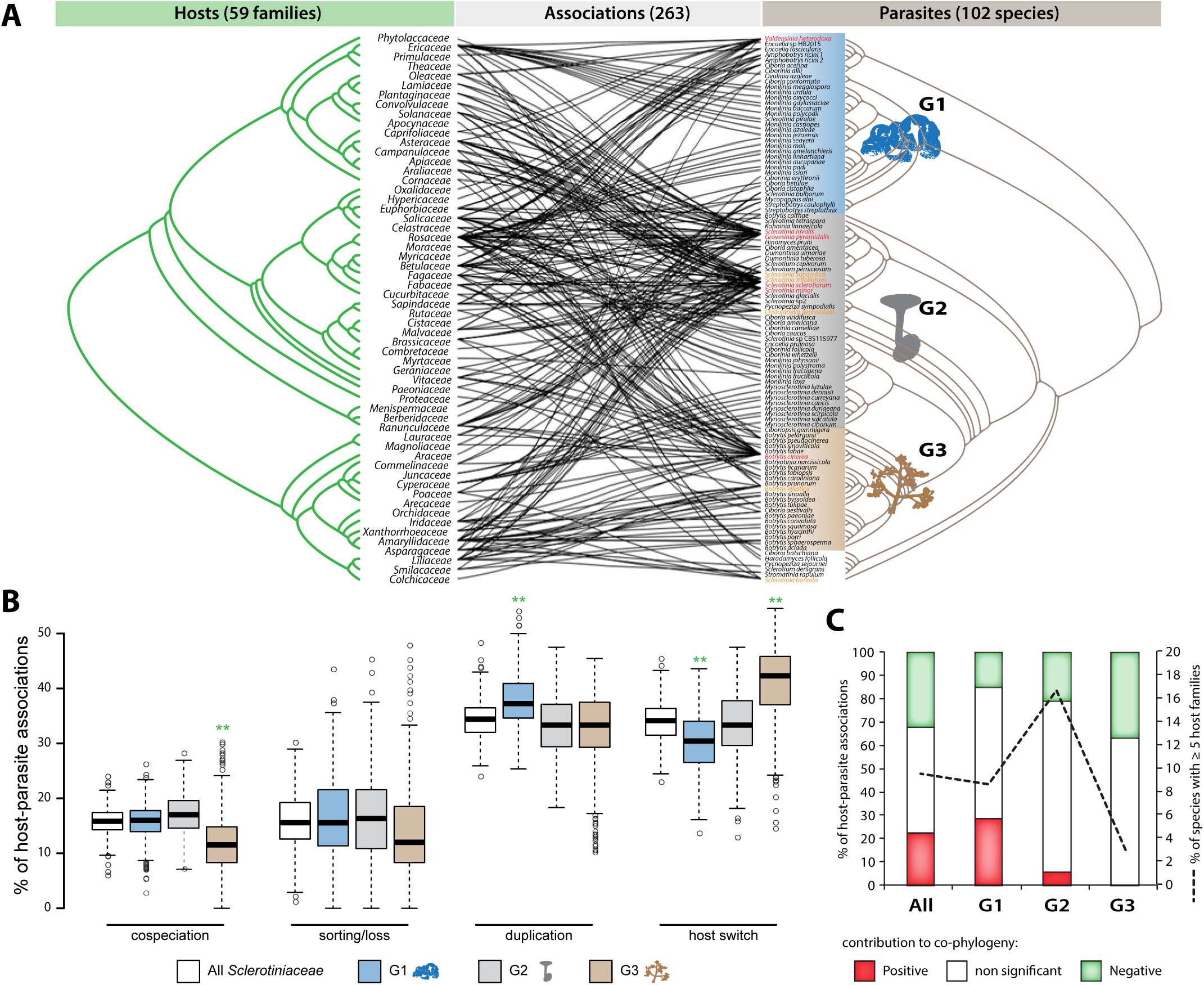
Diversification rate shifts associate with variations in rates of co-speciation, duplication and host switch in *Sclerotiniaceae* fungi. (A) Tanglegram depicting the associations between 102 *Sclerotiniaceae* species and 59 plant families. The three macro-evolutionary regimes are indicated by colored boxes on the *Sclerotiniaceae* tree. Fungal species labels are color-coded as in figure 2. (B) Proportion of co-speciation, sorting/loss, duplication and host switches in *host-Sclerotiniaceae* associations as predicted by CoRe-PA in 1,000 co-phylogeny reconstructions. ** indicate large effect size in a macro-evolutionary compared to the complete set of associations as assessed by Cohen’s d test. **(C)** Proportion of host-*Sclerotiniaceae* associations contributing significantly and positively (likely co-speciation), non significantly and significantly and negatively (likely host switch) to co-phylogeny in PACo analysis. The black dotted line indicates the percentage of broad host range species (5 or more host families) in each group.

## DISCUSSION

Our analyses lead to a model in which the extant diversity of *Sclerotiniaceae* fungi is the result of three macro-evolutionary regimes characterized by distinct diversification rates and host association patterns. Patterns of co-phylogeny decreased from regime G1 to G3, while the frequency of inferred host jump events increased from G1 to G3. Regime G2, which includes the highest proportion of broad host range parasites, showed a frequency of host jumps intermediate between G1 and G3. Our co-phylogeny analyses are consistent with the view that long-term plant-pathogen co-speciation is rare (deVienne et al., 2013). The decrease in host-pathogen co-phylogeny signal from G1 to G3 regime (**Figure 4**) could indicate more frequent true co-speciation events in early diverging *Sclerotiniaceae* species, or could result from host jumps being restricted to closely related hosts for G1 species whereas G2 and G3 species progressively gained the ability to jump to more divergent hosts (deVienne et al., 2007, deVienne et al., 2013). Low diversification rates, notably in regime G2, may have resulted from increased extinction rates. It is conceivable that a high extinction rate of specialist parasites during regime G2 is responsible for the reduced clade diversification rates, while increasing the apparent frequency of emergence of generalist species. This phenomenon, consistent with the view of specialization as “an evolutionary dead-end” (leading to a reduced capacity to diversify), is notably supported in *Tachinidae* parasitic flies and hawkmoths pollinating *Ruellia* plants (Day et al., 2016). An analysis of *Papilionoidea* and *Heliconii* revealed lower rates of diversification for butterfly species feeding on a broad range of plants compared to specialist butterflies (Hardy & Otto, 2014, Day et al., 2016). Consistently, theoretical models of sympatric speciation predict that competition for a narrow range of resources (specialization) to be a strong driver of speciation (Dieckmann & Doebeli, 1999). Similar to plant-feeding insects, the diversity of fungal and oomycete pathogens is considered largely driven by host jumps rather than host specialization that may follow (Hardy & Otto, 2014, Choi & Thines, 2015). Low diversification rates in G2 may also result from a strong increase in diversification rate during the transition from regime G2 to G3. Our ancestral state reconstruction analysis inferred a jump to monocots at the base of regime G3 (**Figure 1**). A recent study on *Hesperiidae* butterflies reported a strong increase in diversification coincident with a switch from dicot-feeders to monocot-feeders (Sahoo et al., 2017). As postulated for *Hesperridae*, the emergence of open grasslands and global temperature decrease may have affected diversification in the *Sclerotiniaceae*.

Competition for resources is likely to lead to speciation if host range expansion is costly (Ackermann et al., 2004). Notably, if different host families are scattered in space and phylogenetically related hosts are clustered, the cost of host range expansion is expected to increase, due to higher costs for dispersal or tradeoffs with other traits (Ackermann et al., 2004). In agreement with this theory, genomic signatures associated with metabolic cost optimization were stronger in generalist than specialist fungal parasites (Badet et al., 2017). Strong climate oscillations during the Cenozoic Era, leading to population isolation through range fragmentations and dispersal events, likely contributed to the radiation of plant lineages (Nyman et al., 2012). For instance, major events in the diversification of *Brassicaceae* plant family coincide with glaciations and arid conditions of the Eocene-Oligocene and the Oligocene-Miocene transitions (Hohmann et al., 2015). In addition, the mid-Miocene (20 to 10 Mya) corresponds to the emergence of the first open grassland habitats in northern Eurasia (Strömberg, 2011). The fragmentation and diversification of plant host populations may have increased the cost for host range expansion in fungal pathogens. A jump to monocots at the base of *Sclerotiniaceae* regime G3 ^~^10 Mya may have favored the conquest of grassland habitats and the rapid diversification of specialist species over the emergence of generalists in this group.

Similar to herbivore diet breadth (Forister et al., 2015), host range in the *Sclerotiniaceae* shows a continuous distribution from specialists to broad host range generalists, with the majority of species being specialists. Mathematical analyses and studies of the host range of insect herbivores suggest that host range expansion could involve the emergence of a “pre-adaptation” followed by the colonization of new hosts (Janz & Nylin, 2008). The existence of such pre-existing enabler traits has been proposed as a facilitator for several shifts in plant species distribution (Donoghue & Edwards, 2014). Notably, the pre-existence of a symbiotic signaling pathway in algae is thought to have facilitated the association of land plants with symbiotic fungi (Delaux et al., 2015). In the plant genus *Hypericum*, increased diversification rates in the Miocene Epoch were likely facilitated by adaptation to colder climates (Nürk et al., 2015). In the bacterial pathogen *Serratia marcescens* and in yeast, adaptation to temperature change was associated with improved tolerance to other stresses (Ketola et al., 2013, Caspeta & Nielsen, 2015). Analysis of the complete predicted proteomes of *S. borealis, S. sclerotiorum* and *Botrytis cinerea* have revealed protein signatures often associated with cold adaptation in the secreted protein of all three species (Badet et al., 2015). Cold tolerance might have been a pre-adaptation that facilitated the emergence of generalist parasites under low diversification rates (regime G2) and a rapid diversification following host jumps on fragmented host populations (regime G3). Indeed, *Sclerotinia borealis*, *S. glacialis*, *S. subarctica* and *S. nivalis*, that diverged during regime G2, are largely restricted to hemiboreal climates (circumboreal region) and can have a lower optimal growth temperature than their sister species (Saito, 1997, Hoshino et al., 2010).

These findings suggest that global climate instability and host diversification in the Cenozoic might have impacted on the diversity of fungal parasites within the *Sclerotiniaceae*. This effect could have been direct, through the emergence of cold adaptation as an enabling trait, or indirect through changes in host population structures and host-parasite association patterns. Knowledge on the dynamics of pathogen evolution increases the understanding of the complex interplay between host, pathogen and environment governing the dynamics of disease epidemics. These evolutionary principles are useful for the design of disease management strategies (Vander Wal et al., 2014) and provide new insights into the factors that influenced the diversity of extant fungal parasite species.

## ACKNOWLEDGEMENTS

This work was supported by grants from the European Research Council (ERC-StG-336808 to S.R.), the French Laboratory of Excellence project TULIP (ANR-10-LABX-41; ANR-11-IDEX-0002-02) and a "New Frontiers" grant of the Laboratory of Excellence project TULIP. We thank the TULIP international advisory board and the TULIP community for stimulating discussions.

## Authors contributions

ON: collected data, performed analyses, writing original draft and revised manuscript

AB: performed analyses, writing revised manuscript

AT: collected data, revised manuscript

JPC: funding acquisition, collected data, revised manuscript

SR: Supervision, funding acquisition, project administration, writing original draft and revised manuscript

## DATA ACCESSIBILITY

- Species identifiers: Genbank accessions provided in Supplementary Table 1
- DNA sequences and alignments : provided as supplementary file 2
- Phylogenetic trees: provided as supplementary file 1, 3, 4 and 5

## SUPPLEMENTARY FILES

**Supplementary Figure 1.**
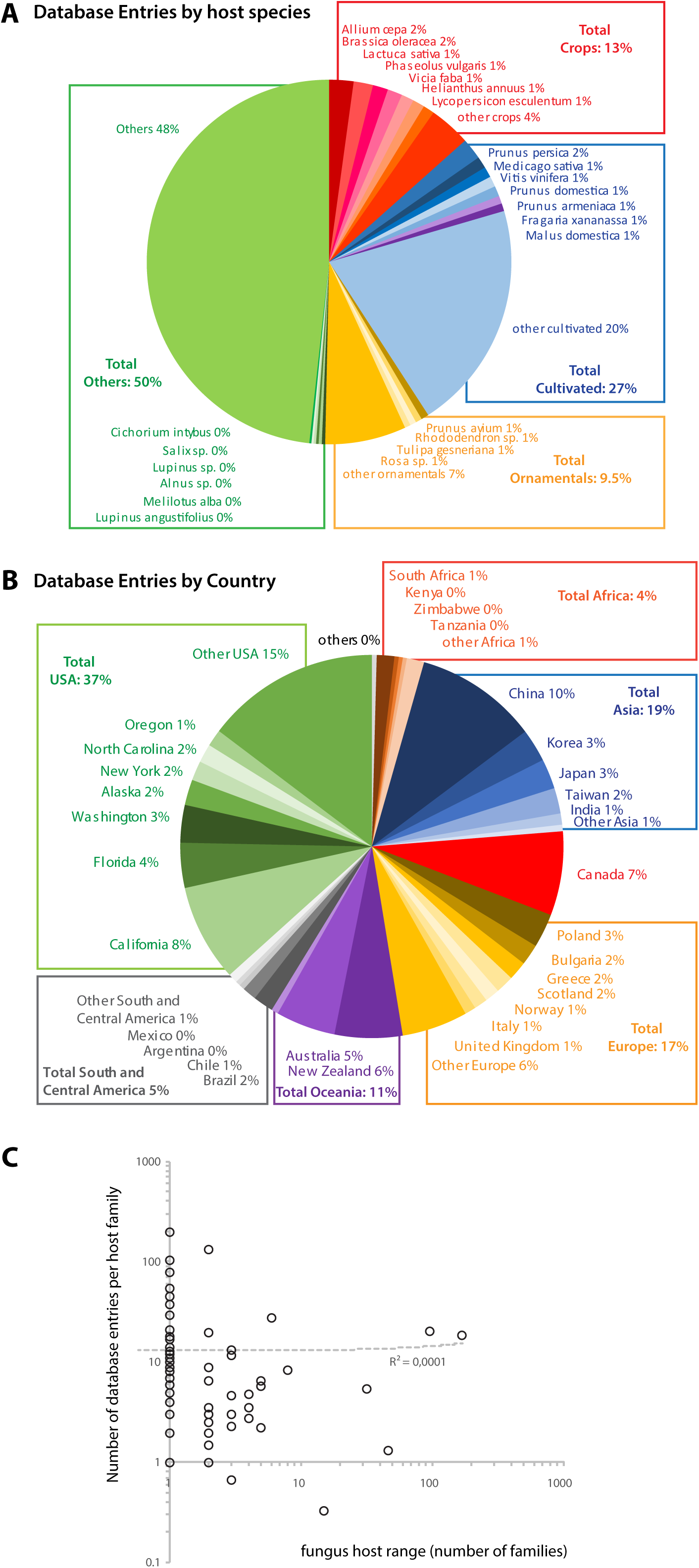
We analyzed the 7101 records from the fungus-host distribution database that served as a basis for our study on the *Sclerotiniaceae.* Host species were classified as crops according to the USDA Natural Resources Conservation Service database (https://plants.usda.gov/npk/main), other cultivated plants, ornamental plants and “others”, including largely wild plant species. We found that 13% of the Fungus-Host database entries related to crops, 20% to cultivated plants and 9.5% to ornamentals (A). A total of 37% of the Fungus-Host database entries were from the USA, 17% from Europe, leaving a total of 46% of the entries that were neither from the USA or Europe (B). These distributions do not reveal a strong bias towards crops or USA/Europe records in the records used for this analysis. (C) Relationship between fungal pathogen host range and the number of records in the database for each plant family. Dotted line shows linear regression of the data.

**Supplementary Figure 2.**
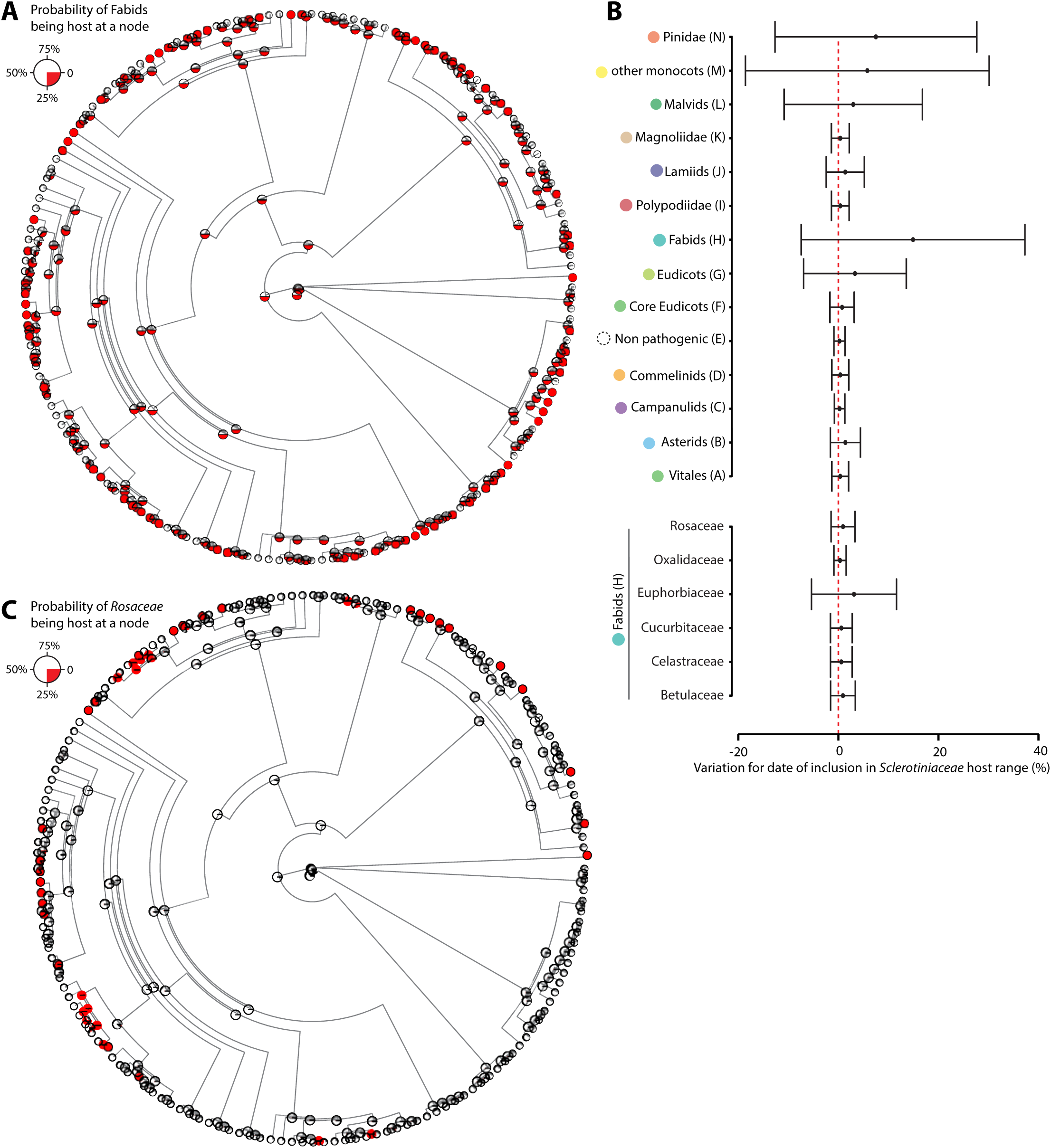
Reconstruction of ancestral host range by the re-rooting method under an entity-relationship model and robustness of the results to tree pruning. (A) Posterior probability for the presence of Fabids in *Sclerotiniaceae* at each internal node (red), Fabids being the most likely ancestral host group of the *Sclerotiniaceae* family in S-DIVA and S-DEC analyses. (B) Sensitivity to tree pruning for the date of inclusion of plant groups into *Sclerotiniaceae* pathogen host range. The graph shows the % of variation for the earliest date of inclusion into *Sclerotiniaceae* pathogen host range upon pruning of 10% of the phylogenetic tree 100 times randomly. Error bars show 95% confidence intervals. (C) Posterior probability for the presence of *Rosaceae* (a family in the Fabids) in *Sclerotiniaceae* host groups at each internal node (red).

**Supplementary Figure 3.**
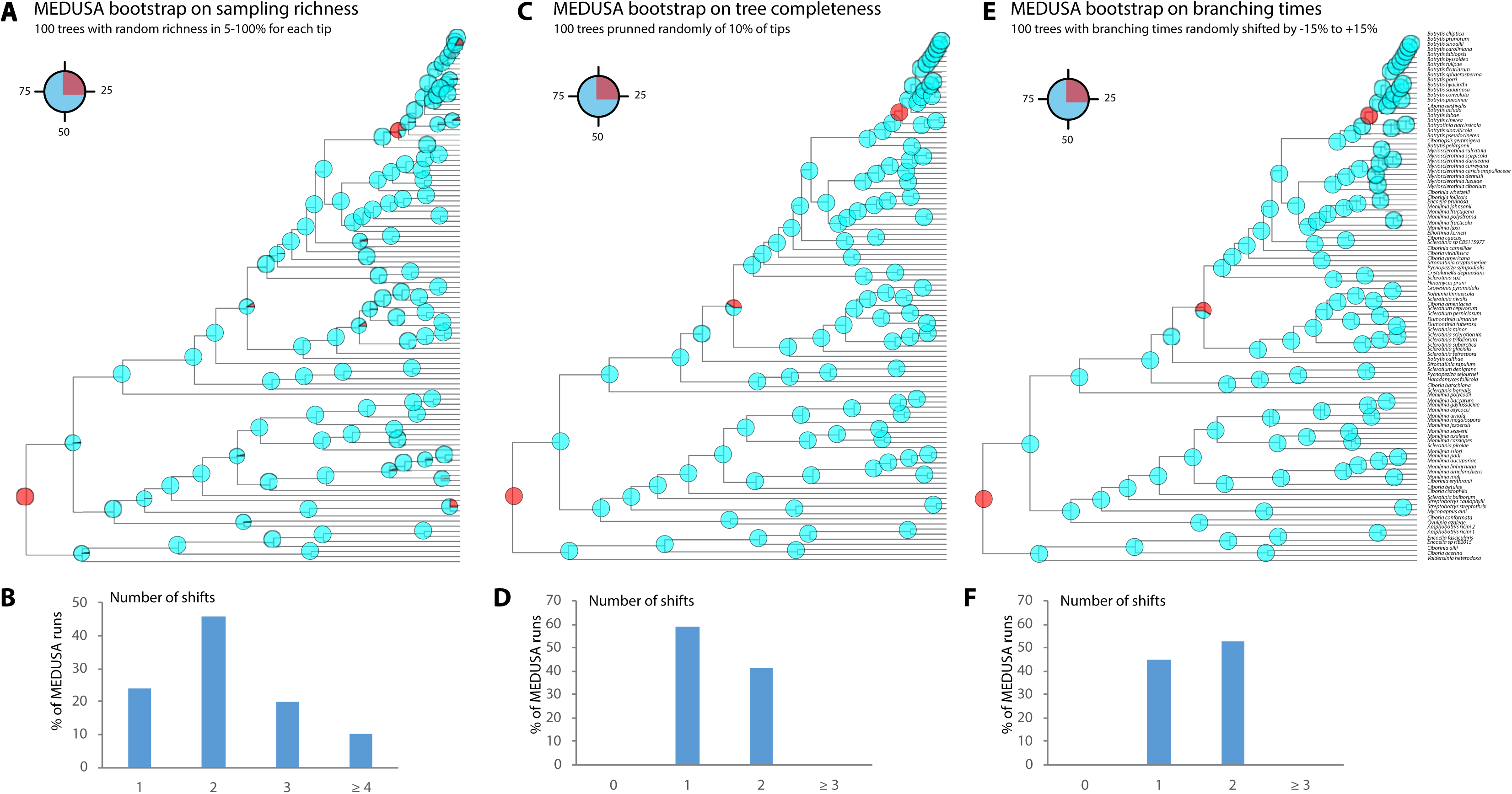
Assessment of the robustness of diversification rate shifts predicted by MEDUSA to sampling richness (A-B), tree completeness (C-D) and dating accuracy (E-F). The figure shows the frequency of detection of a diversification rate shift at every internal node of the tree in 100 bootstrap MEDUSA analyses (A, C and E), as well as the total number of shifts predicted in each of the 100 bootstrap analyses (B, D and F).

**Supplementary Figure 4.**
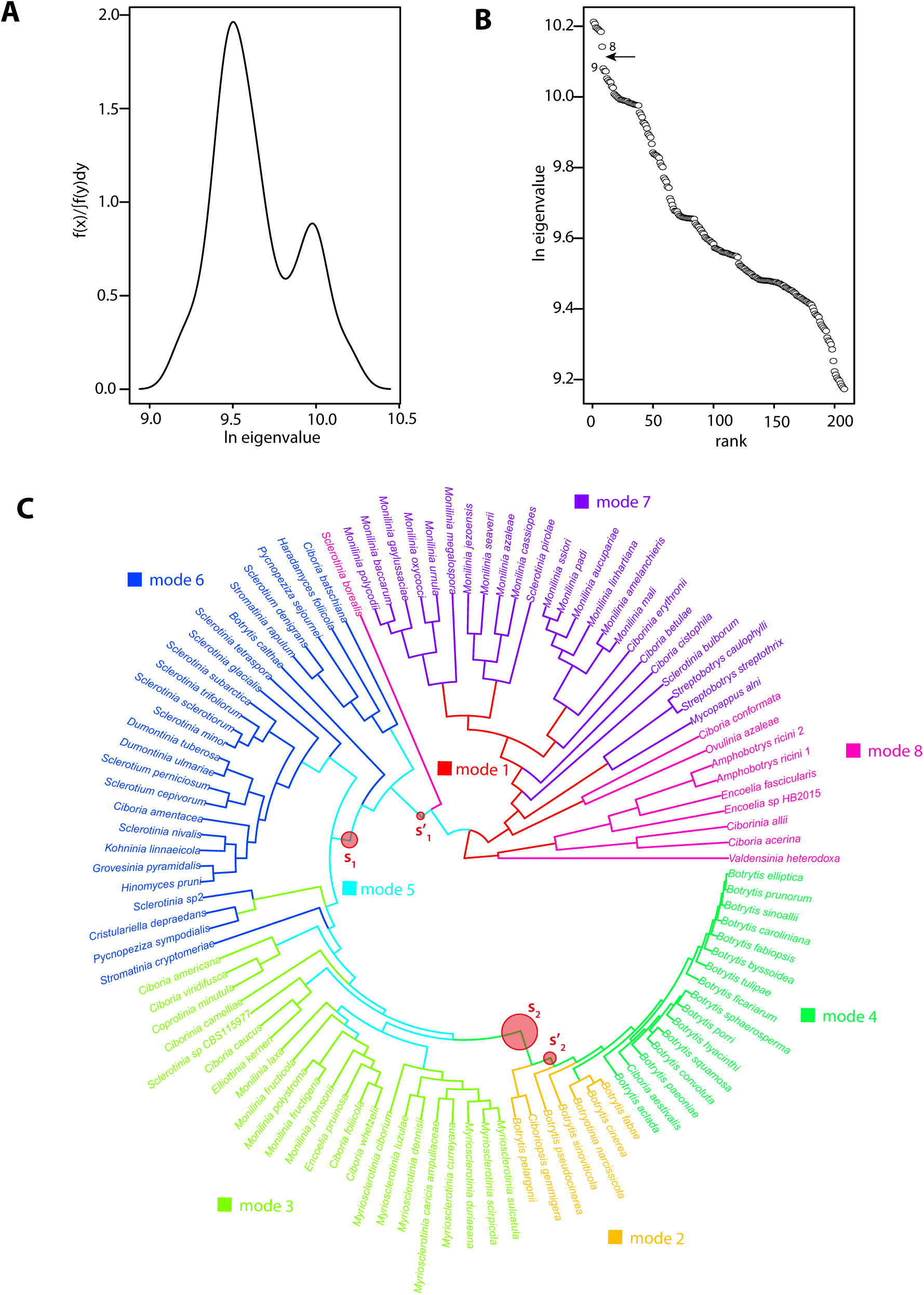
Diversification rates analysis with R-PANDA. (A) Spectral density plot of the *Sclerotiniaceae.* (B) Eigenvalues ranked in descending order for the *Sclerotiniaceae*, the eigengap between values 8 and 9 is shown by an arrow and indicates eight modalities in this phylogeny. (C) Location of the eight modalities on the *Sclerotiniaceae* phylogeny, indicated by different colors. The major rate shifts identified by BAMM are shown as circles on the phylogeny and delimit the shift between modes 1/7/8 to modes 5/6/3 and finally modes 2/4.

**Supplementary figure 5.**
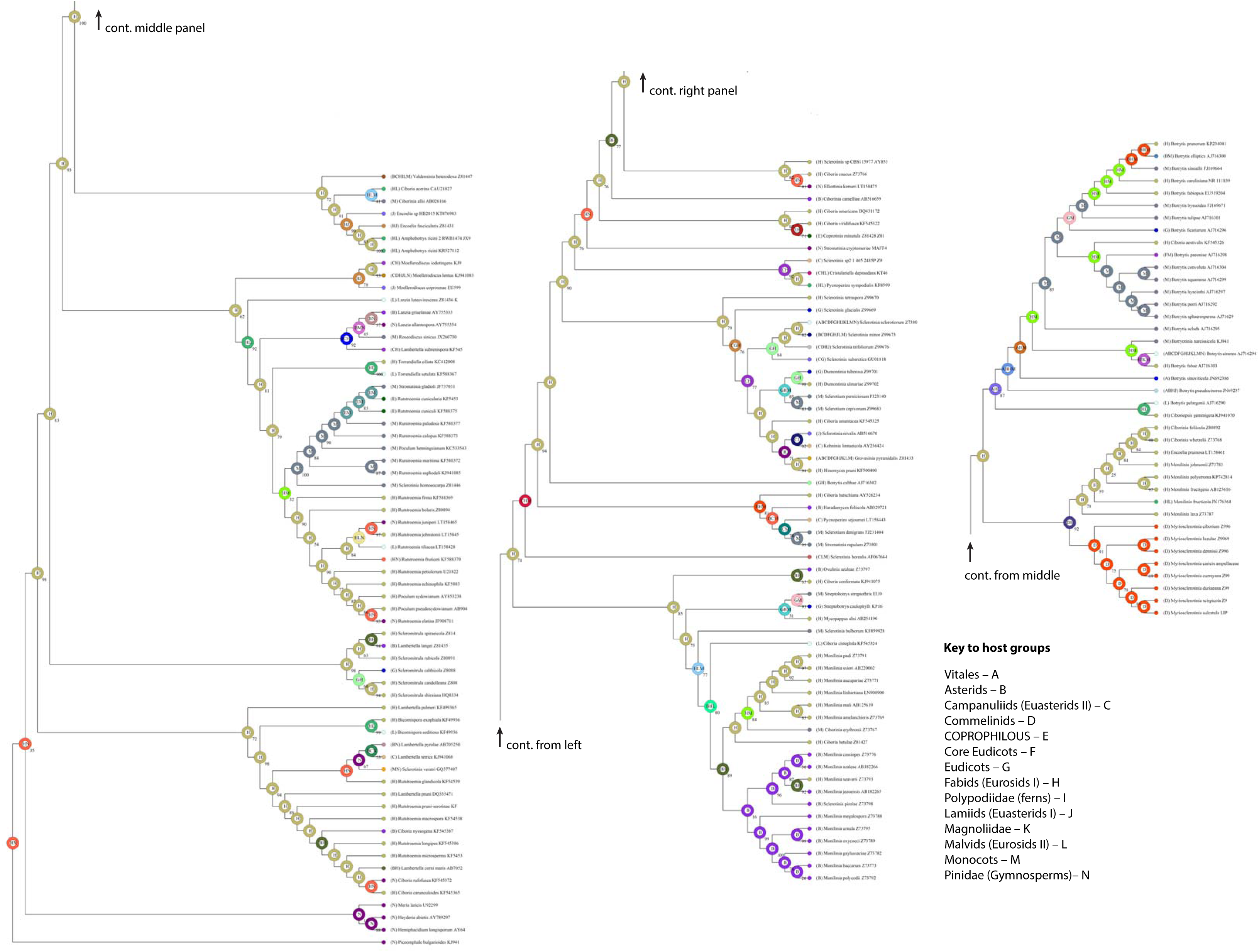
RASP phylogram showing extant and reconstructed ancestral host range for 161 *Sclerotiniaceae* and *Rutstroemiaceae* species. The ancestral reconstruction shows the probabilities associated with the optimal distribution at major nodes. It was generated using the S-DIVA method and the phylogenetic tree provided in supplementary file 2 as a template. The code for host groups is as follows: Vitales (A), Asterids (B), Campanuliids (C), Commelinids (D), coprophilous (E), core Eudicots (F), Eudicots (G), Fabids (H), Polypodiidae (I), Lamiids (J), Magnoloidae (K), Malvids (L), Monocotyledones (M), Pinidae (N).

**Supplementary Figure 6.**
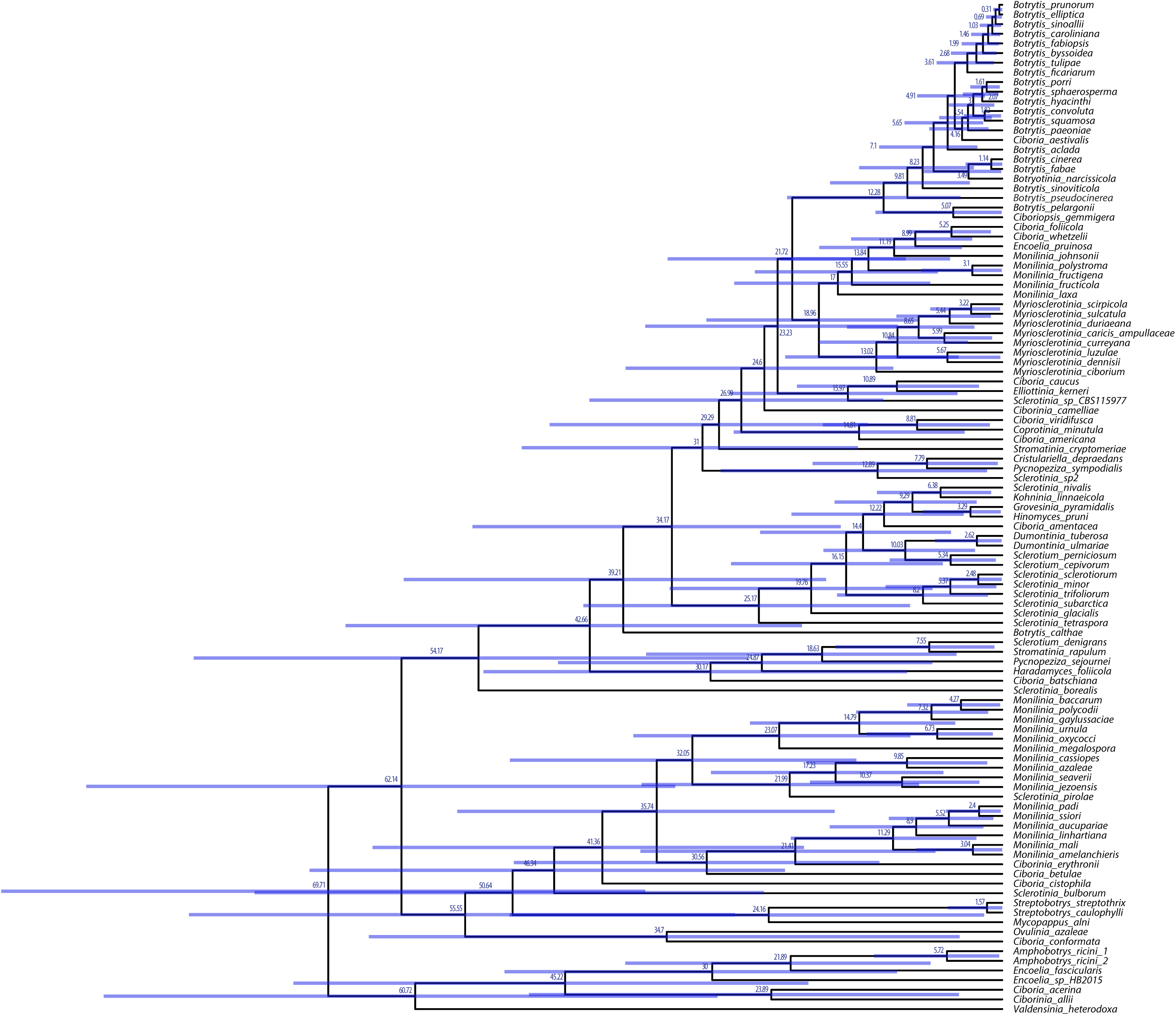
Chronotree of the 105 *Sclerotiniaceae* species generated with BEAST. Estimated mean age of all intermediate nodes is indicated (in million years), with bars showing the 95% confidence interval of the highest posterior density (HPD).

**Supplementary Figure 7.**
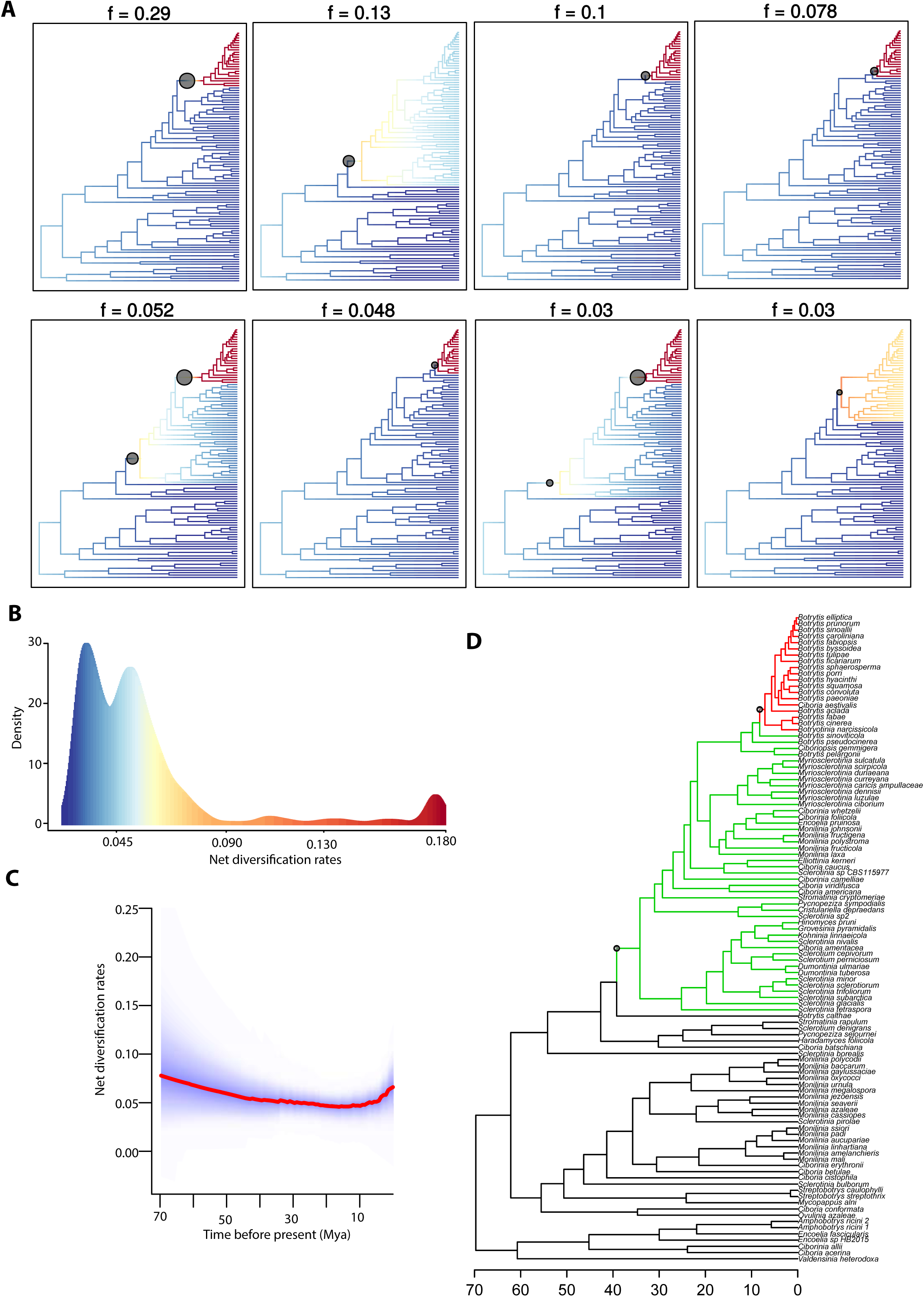
Diversification rates analysis with BAMM and Medusa. (A) The eight most probable shift configurations in the 95% credible set of shift configurations predicted by BAMM with the posterior probability of each configuration indicated on top. (B) Density distribution of net diversification rates calculated by BAMM. The three peaks of density reflect three macro-evolutionary regimes in the *Sclerotiniaceae.* (C) Net diversification rates as a function of time calculated by BAMM through the complete *Sclerotiniaceae* phylogeny (105 species), showing a first a relatively sharp decrease (70 to 55 Mya), followed by a slow decrease (55 to 17 Mya) and a recent increase (17 Mya to present) in diversification rates. (D) Diversification rate shifts identified by MEDUSA.

**Supplementary File 1.** Phylogenetic tree of host plant families used for co-phylogenetic analyses in this work (newick format).

**Supplementary File 2.** Curated multiple ITS sequence alignment for 200 Leotiomycete species, including 105 *Sclerotiniaceae* and 56 *Rutstroemiaceae* species. This alignment includes 797 informative sites and was used to generate the phylogenetic tree shown in Figure 1 and downstream analyses (fasta format).

**Supplementary File 3.** Phylogenetic tree of 200 Leotiomycete species including 161 *Sclerotiniaceae* and *Rutstroemiaceae* species used in Figure 1, obtained by maximum likelihood approach and featuring SH-aLRT branch support (newick format).

**Supplementary File 4**. Same phylogenetic tree as in supplementary file 3 including bootstrap from 100 replicates as branch support (newick format).

**Supplementary File 5.** Time calibrated phylogenetic tree of the 105 *Sclerotiniaceae* species used in Figure 2.

**Supplementary File 6.** List of host-parasite associations tested for co-phylogenetic analyses.

**Supplementary Table 1.** List of *Sclerotiniaceae* and *Rutstroemiaceae* species used for phylogenetic analysis and their corresponding host range. 1 refers to position in the tree shown in Figure 1; 2 refers to the code used in RASP analysis (Sup. Figure 5). NA, not applicable; Rutst., *Rutstroemiaceae;* Sclero. *Sclerotiniaceae*.

## REFERENCES

Ackermann M, Doebeli M, Gomulkiewicz R, 2004. Evolution of niche width and adaptive diversification. Evolution 58, 2599–612.

Alfaro ME, Santini F, Brock C, et al., 2009. Nine exceptional radiations plus high turnover explain species diversity in jawed vertebrates. Proceedings of the National Academy of Sciences 106, 13410–4.

Andrew M, Barua R, Short SM, Kohn LM, 2012. Evidence for a Common Toolbox Based on Necrotrophy in a Fungal Lineage Spanning Necrotrophs, Biotrophs, Endophytes, Host Generalists and Specialists. PLoS One 7, e29943.

Anisimova M, Gil M, Dufayard J-F, Dessimoz C, Gascuel O, 2011. Survey of branch support methods demonstrates accuracy, power, and robustness of fast likelihood-based approximation schemes. Systematic biology, syr041.

Badet T, Peyraud R, Mbengue M, et al., 2017. Codon optimization underpins generalist parasitism in fungi. Elife 6, e22472.

Badet T, Peyraud R, Raffaele S, 2015. Common protein sequence signatures associate with *Sclerotinia borealis* lifestyle and secretion in fungal pathogens of the *Sclerotiniaceae*. Front Plant Sci 6, 776.

Balbuena JA, Míguez-Lozano R, Blasco-Costa I, 2013. PACo: a novel procrustes application to cophylogenetic analysis. PLoS One 8, e61048.

Barrett LG, Kniskern JM, Bodenhausen N, Zhang W, Bergelson J, 2009. Continua of specificity and virulence in plant host–pathogen interactions: causes and consequences. New Phytologist 183, 513–29.

Beimforde C, Feldberg K, Nylinder S, et al., 2014. Estimating the Phanerozoic history of the Ascomycota lineages: combining fossil and molecular data. Molecular phylogenetics and evolution 78, 386–98.

Boland G, Hall R, 1994. Index of plant hosts of Sclerotinia sclerotiorum. Canadian Journal of Plant Pathology 16, 93–108.

Bolton MD, Thomma BPHJ, Nelson BD, 2006. Sclerotinia sclerotiorum (Lib.) de Bary: biology and molecular traits of a cosmopolitan pathogen. Molecular Plant Pathology 7, 1–16.

Caspeta L, Nielsen J, 2015. Thermotolerant yeast strains adapted by laboratory evolution show trade-off at ancestral temperatures and preadaptation to other stresses. MBio 6, e00431-15.

Chappell TM, Rausher MD, 2016. Evolution of host range in Coleosporium ipomoeae, a plant pathogen with multiple hosts. Proceedings of the National Academy of Sciences 113, 5346–51.

Charleston M, Robertson D, 2002. Preferential host switching by primate lentiviruses can account for phylogenetic similarity with the primate phylogeny. Systematic biology 51, 528–35.

Choi Y-J, Thines M, 2015. Host jumps and radiation, not co-divergence drives diversification of obligate pathogens. A case study in downy mildews and Asteraceae. PLoS One 10, e0133655.

Clarke JT, Warnock R, Donoghue PC, 2011. Establishing a time-scale for plant evolution. New Phytologist 192, 266–301.

Clarkson JP, Carter H, Coventry E, 2010. First report of Sclerotinia subarctica nom. prov.(Sclerotinia species 1) in the UK on Ranunculus acris. Plant pathology 59, 1173-.

Conow C, Fielder D, Ovadia Y, Libeskind-Hadas R, 2010. Jane: a new tool for the cophylogeny reconstruction problem. Algorithms for Molecular Biology 5, 16.

Day EH, Hua X, Bromham L, 2016. Is specialization an evolutionary dead end? Testing for differences in speciation, extinction and trait transition rates across diverse phylogenies of specialists and generalists. Journal of evolutionary biology 29, 1257–67.

Dean R, Van Kan JaL, Pretorius ZA, et al., 2012. The Top 10 fungal pathogens in molecular plant pathology. Molecular Plant Pathology 13, 414–30.

Delaux P-M, Radhakrishnan GV, Jayaraman D, et al., 2015. Algal ancestor of land plants was preadapted for symbiosis. Proceedings of the National Academy of Sciences 112, 13390–5.

Denton-Giles M, Bradshaw RE, Dijkwel PP, 2013. Ciborinia camelliae (Sclerotiniaceae) induces variable plant resistance responses in selected species of Camellia. Phytopathology 103, 725–32.

Devienne D, Giraud T, Shykoff J, 2007. When can host shifts produce congruent host and parasite phylogenies? A simulation approach. Journal of evolutionary biology 20, 1428–38.

Devienne D, Refrégier G, López-Villavicencio M, Tellier A, Hood M, Giraud T, 2013. Cospeciation vs host-shift speciation: methods for testing, evidence from natural associations and relation to coevolution. New Phytologist 198, 347–85.

Dieckmann U, Doebeli M, 1999. On the origin of species by sympatric speciation. Nature 400, 354–7.

Dong S, Raffaele S, Kamoun S, 2015. The two-speed genomes of filamentous pathogens: waltz with plants. Curr Opin Genet Dev 35, 57–65.

Donoghue MJ, Edwards EJ, 2014. Biome shifts and niche evolution in plants. Annual Review of Ecology, Evolution, and Systematics 45, 547–72.

Drummond AJ, Suchard MA, Xie D, Rambaut A, 2012a. Bayesian phylogenetics with BEAUti and the BEAST 1.7. Molecular biology and Evolution 29, 1969–73.

Drummond CS, Eastwood RJ, Miotto ST, Hughes CE, 2012b. Multiple continental radiations and correlates of diversification in Lupinus (Leguminosae): testing for key innovation with incomplete taxon sampling. Systematic biology 61, 443–60.

Elliott ME, 1967. Rutstroemia cuniculi, a coprophilous species of the Sclerotiniaceae. Canadian Journal of Botany 45, 521–4.

Farr DF, Rossman AY, 2016. Fungal Databases, Systematic Mycology and Microbiology Laboratory, ARS, USDA.

Forister ML, Novotny V, Panorska AK, et al., 2015. The global distribution of diet breadth in insect herbivores. Proceedings of the National Academy of Sciences 112, 442–7.

Futuyma DJ, Moreno G, 1988. The evolution of ecological specialization. Annual Review of Ecology and Systematics, 207–33.

Gouy M, Guindon S, Gascuel O, 2010. SeaView version 4: a multiplatform graphical user interface for sequence alignment and phylogenetic tree building. Molecular biology and Evolution 27, 221–4.

Graf F, Schumacher T, 1995. Sclerotinia glacialis sp. nov., from the alpine zone of Switzerland. Mycological Research 99, 113–7.

Guindon S, Dufayard J-F, Lefort V, Anisimova M, Hordijk W, Gascuel O, 2010. New algorithms and methods to estimate maximum-likelihood phylogenies: assessing the performance of PhyML 3.0. Systematic biology 59, 307–21.

Haldane JBS, 1951. Everything has a History. London: Routledge.

Hamm CA, Fordyce JA, 2015. Patterns of host plant utilization and diversification in the brush-footed butterflies. Evolution 69, 589–601.

Hardy NB, Otto SP, 2014. Specialization and generalization in the diversification of phytophagous insects: tests of the musical chairs and oscillation hypotheses. Proceedings of the Royal Society of London B: Biological Sciences 281, 2013–2960.

Hawksworth DL, Luecking R, 2017. Fungal Diversity Revisited: 2.2 to 3.8 Million Species. Microbiology spectrum 5.

Hedges SB, Marin J, Suleski M, Paymer M, Kumar S, 2015. Tree of life reveals clock-like speciation and diversification. Molecular biology and Evolution 32, 835–45.

Hohmann N, Wolf EM, Lysak MA, Koch MA, 2015. A time-calibrated road map of Brassicaceae species radiation and evolutionary history. The Plant Cell 27, 2770–84.

Holst-Jensen A, Kohn L, Jakobsen K, Schumacher T, 1997. Molecular phylogeny and evolution of Monilinia (Sclerotiniaceae) based on coding and noncoding rDNA sequences. American journal of botany 84, 686-.

Holst-Jensen A, Vrålstad T, Schumacher T, 2004. Kohninia linnaeicola, a new genus and species of the Sclerotiniaceae pathogenic to Linnaea borealis. Mycologia 96, 135–42.

Holst-Jensen A, Vaage M, Schumacher T, 1998. An approximation to the phylogeny of Sclerotinia and related genera. Nordic Journal of Botany 18, 705–19.

Hoshino T, Terami F, Tkachenko OB, Tojo M, Matsumoto N, 2010. Mycelial growth of the snow mold fungus, Sclerotinia borealis, improved at low water potentials: an adaption to frozen environment. Mycoscience 51, 98–103.

Hu X, Xiao G, Zheng P, et al., 2014. Trajectory and genomic determinants of fungal-pathogen speciation and host adaptation. Proceedings of the National Academy of Sciences 111, 16796–801.

Janz N, Nylin S, 2008. The oscillation hypothesis of host-plant range and speciation. In: Tilmon K, ed. Specialization, speciation, and radiation: the evolutionary biology of herbivorous insects. Univ. of California Press, Berkeley, CA., 203–15.

Janz N, Nylin S, Wahlberg N, 2006. Diversity begets diversity: host expansions and the diversification of plant-feeding insects. BMC evolutionary biology 6, 4.

Johnson KP, Malenke JR, Clayton DH, 2009. Competition promotes the evolution of host generalists in obligate parasites. Proceedings of the Royal Society of London B: Biological Sciences 276, 3921–6.

Joshi A, Thompson JN, 1995. Trade-offs and the evolution of host specialization. Evolutionary Ecology 9, 82–92.

Katoh K, Standley DM, 2013. MAFFT multiple sequence alignment software version 7: improvements in performance and usability. Molecular biology and Evolution 30, 772–80.

Ketola T, Mikonranta L, Zhang J, et al., 2013. Fluctuating temperature leads to evolution of thermal generalism and preadaptation to novel environments. Evolution 67, 2936–44.

Lefort V, Desper R, Gascuel O, 2015. FastME 2.0: a comprehensive, accurate, and fast distance-based phylogeny inference program. Molecular biology and Evolution 32, 2798–800.

Lewitus E, Morlon H, 2015. Characterizing and comparing phylogenies from their Laplacian spectrum. Systematic biology 65, 495–507.

Liao J, Huang H, Meusnier I, et al., 2016. Pathogen effectors and plant immunity determine specialization of the blast fungus to rice subspecies. Elife 5, e19377.

Lorenzini M, Zapparoli G, 2016. Description of a taxonomically undefined Sclerotiniaceae strain from withered rotten-grapes. Antonie van Leeuwenhoek 109, 197–205.

Mbengue M, Navaud O, Peyraud R, et al., 2016. Emerging trends in molecular interactions between plants and the broad host range fungal pathogens Botrytis cinerea and Sclerotinia sclerotiorum. Frontiers in plant science 7.

Mcmanus P, Best V, Voland R, 1999. Infection of cranberry flowers by Monilinia oxycocci and evaluation of cultivars for resistance to cottonball. Phytopathology 89, 1127–30.

Melzer M, Smith E, Boland G, 1997. Index of plant hosts of Sclerotinia minor. Canadian Journal of Plant Pathology 19, 272–80.

Merkle D, Middendorf M, Wieseke N, 2010. A parameter-adaptive dynamic programming approach for inferring cophylogenies. BMC Bioinformatics 11, S60.

Moran NA, 1988. The evolution of host-plant alternation in aphids: evidence for specialization as a dead end. The American naturalist 132, 681–706.

Morlon H, Lewitus E, Condamine FL, Manceau M, Clavel J, Drury J, 2016. RPANDA: an R package for macroevolutionary analyses on phylogenetic trees. Methods in Ecology and Evolution.

Nürk NM, Uribe-Convers S, Gehrke B, Tank DC, Blattner FR, 2015. Oligocene niche shift, Miocene diversification–cold tolerance and accelerated speciation rates in the St. John's Worts (Hypericum, Hypericaceae). BMC evolutionary biology 15, 80.

Nyman T, Linder HP, Peña C, Malm T, Wahlberg N, 2012. Climate-driven diversity dynamics in plants and plant-feeding insects. Ecology letters 15, 889–98.

Paradis E, Claude J, Strimmer K, 2004. APE: analyses of phylogenetics and evolution in R language. Bioinformatics 20, 289–90.

Poulin R, Keeney DB, 2008. Host specificity under molecular and experimental scrutiny. Trends in parasitology 24, 24–8.

Prieto M, Wedin M, 2013. Dating the diversification of the major lineages of Ascomycota (Fungi). PLoS One 8, e65576.

Qian H, Zhang J, 2014. Using an updated time-calibrated family-level phylogeny of seed plants to test for non-random patterns of life forms across the phylogeny. Journal of systematics and evolution 52, 423–30.

Rabosky DL, 2014. Automatic detection of key innovations, rate shifts, and diversity-dependence on phylogenetic trees. PLoS One 9, e89543.

Rabosky DL, Grundler M, Anderson C, et al., 2014. BAMMtools: an R package for the analysis of evolutionary dynamics on phylogenetic trees. Methods in Ecology and Evolution 5, 701–7.

Rabosky DL, Mitchell JS, Chang J, 2017. Is BAMM flawed? Theoretical and practical concerns in the analysis of multi-rate diversification models. Systematic biology.

Rabosky DL, Santini F, Eastman J, et al., 2013. Rates of speciation and morphological evolution are correlated across the largest vertebrate radiation. Nature Communications 4.

Razo-Mendivil U, De Leon GP-P, 2011. Testing the evolutionary and biogeographical history of Glypthelmins (Digenea: Plagiorchiida), a parasite of anurans, through a simultaneous analysis of molecular and morphological data. Molecular phylogenetics and evolution 59, 331–41.

Revell LJ, 2012. phytools: an R package for phylogenetic comparative biology (and other things). Methods in Ecology and Evolution 3, 217–23.

Ronquist F, Teslenko M, Van Der Mark P, et al., 2012. MrBayes 3.2: efficient Bayesian phylogenetic inference and model choice across a large model space. Systematic biology 61, 539–42.

Sahoo RK, Warren AD, Collins SC, Kodandaramaiah U, 2017. Hostplant change and paleoclimatic events explain diversification shifts in skipper butterflies (Family: Hesperiidae). BMC evolutionary biology 17, 174.

Saito I, 1997. Sclerotinia nivalis, sp. nov., the pathogen of snow mold of herbaceous dicots in northern Japan. Mycoscience 38, 227.

Schliep KP, 2010. phangorn: phylogenetic analysis in R. Bioinformatics, btq706.

Schoch CL, Seifert KA, Huhndorf S, et al., 2012. Nuclear ribosomal internal transcribed spacer (ITS) region as a universal DNA barcode marker for Fungi. Proceedings of the National Academy of Sciences 109, 6241–6.

Schumacher T, Kohn LM, 1985. A monographic revision of the genus Myriosclerotinia. Canadian Journal of Botany 63, 1610–40.

Smith SA, Beaulieu JM, Donoghue MJ, 2010. An uncorrelated relaxed-clock analysis suggests an earlier origin for flowering plants. Proceedings of the National Academy of Sciences 107, 5897–902.

Spanu PD, Abbott JC, Amselem J, et al., 2010. Genome expansion and gene loss in powdery mildew fungi reveal tradeoffs in extreme parasitism. Science 330, 1543.

Staats M, Van Baarlen P, Van Kan JA, 2005. Molecular phylogeny of the plant pathogenic genus Botrytis and the evolution of host specificity. Molecular biology and Evolution 22, 333–46.

Strömberg CA, 2011. Evolution of grasses and grassland ecosystems. Annual Review of Earth and Planetary Sciences 39, 517–44.

Thines M, Choi Y-J, 2015. Evolution, diversity, and taxonomy of the Peronosporaceae, with focus on the genus Peronospora. Phytopathology 106, 6–18.

Vander Wal E, Garant D, Calmé S, et al., 2014. Applying evolutionary concepts to wildlife disease ecology and management. Evolutionary applications 7, 856–68.

Visser B, Le Lann C, Den Blanken FJ, Harvey JA, Van Alphen JJ, Ellers J, 2010. Loss of lipid synthesis as an evolutionary consequence of a parasitic lifestyle. Proceedings of the National Academy of Sciences 107, 8677–82.

Walker A-S, Gautier A, Confais J, et al., 2011. Botrytis pseudocinerea, a new cryptic species causing gray mold in French vineyards in sympatry with Botrytis cinerea. Phytopathology 101, 1433–45.

Woolhouse ME, Gowtage-Sequeria S, 2005. Host range and emerging and reemerging pathogens. Emerg Infect Dis 11, 1842–47.

Woolhouse ME, Taylor LH, Haydon DT, 2001. Population biology of multihost pathogens. Science 292, 1109–12.

Yang Z, Kumar S, Nei M, 1995. A new method of inference of ancestral nucleotide and amino acid sequences. Genetics 141, 1641–50.

Yu Y, Harris AJ, Blair C, He X, 2015. RASP (Reconstruct Ancestral State in Phylogenies): a tool for historical biogeography. Molecular phylogenetics and evolution 87, 46–9.

Zachos J, Pagani M, Sloan L, Thomas E, Billups K, 2001. Trends, rhythms, and aberrations in global climate 65 Ma to present. Science 292, 686–93.

